## Supplementary informations for "NK Cell Exhaustion in Wilson’s Disease Revealed by Single-cell RNA Sequencing Predicts the Prognosis of Cholecystitis"

Figure S1

a.

**Clinical characteristics of the patients at baseline.**

| Cohort 1 | | | Cohort 2 | | | Cohort 3 | | |
|---|---|---|---|---|---|---|---|---|
| | WD (n=92) | HBV (n=95) | | WD (n=618) | HBV (n=370) | | Cholecystitis & WD (n=15) | Cholecystitis (n=17) |
| Age - yr | 28.62±9.67 | 55.76±15.88 | Age - yr | 31.60±11.33 | 49.85±13.77 | Age - yr | 32.87±10.72 | 52.12±16.64 |
| Male sex – no. (%) | 63 (68.48%) | 65 (68.42%) | Male sex – no. (%) | 357 (57.77%) | 254 (68.64%) | Male sex – no. (%) | 9 (60.00%) | 8 (47.06%) |

WD, Wilson disease; HBV, Hepatitis B virus.

b.

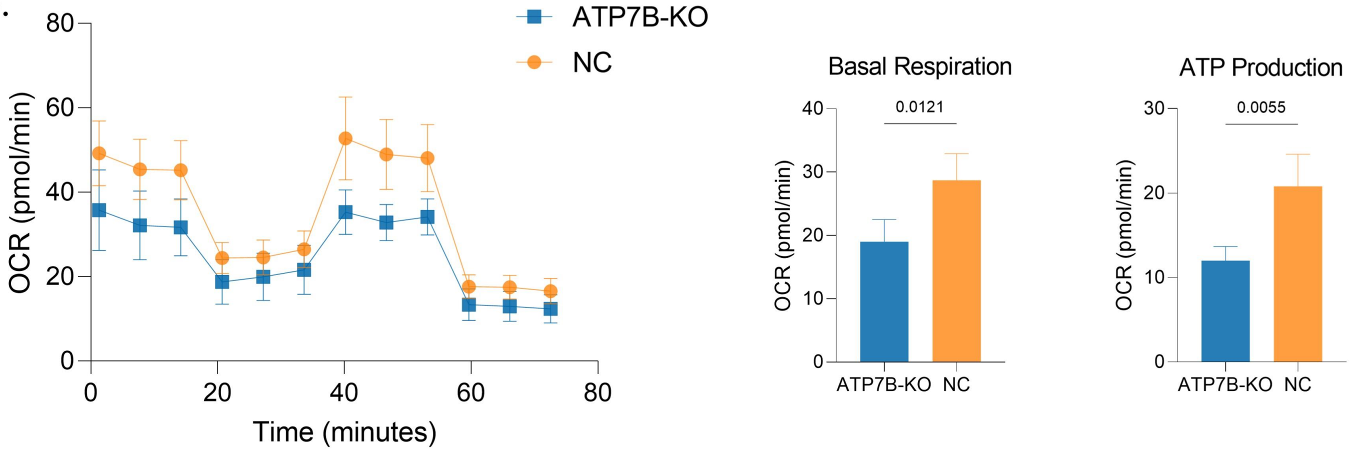

c.

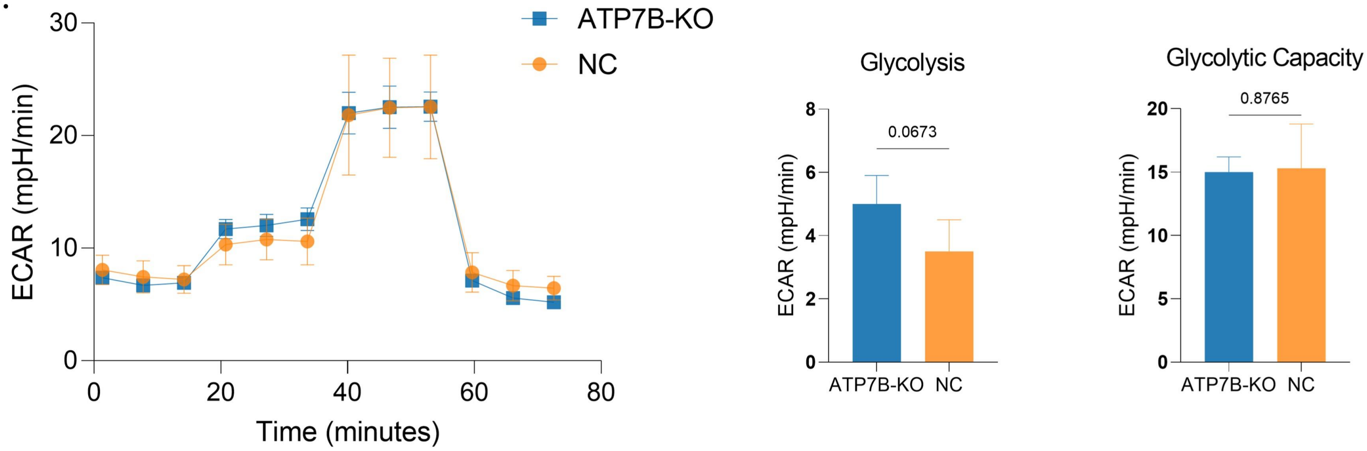

Figure S2

a.

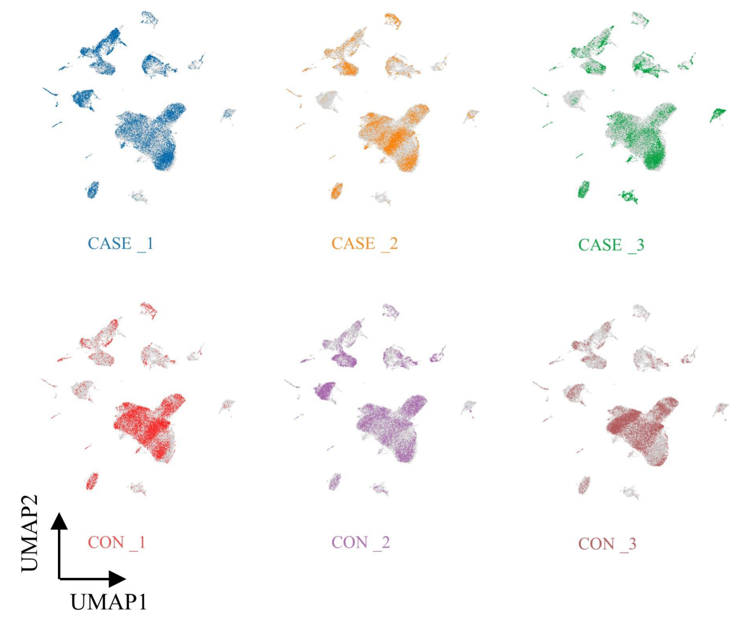

c.

**Cell marker genes for annotation of main cell types.**

| Celltype | Markers |
|---|---|
| Hepatocytes | ALB,TTR,APOA1 |
| Cholangiocytes | KRT19,KRT7,CFTR |
| Endothelial cells | PECAM1,VWF,CDH5 |
| Hepatic stellate cells | PDGFRB,ACTA2,RGS5 |
| Proliferating cells | MKI67,TOP2A,STMN1 |
| B cells | CD79A,MS4A1,CD19 |
| Plasma cells | JCHAIN,CD79A,MZB1 |
| T and NK cells | CD3D,CD3E,NKG7 |
| Neutrophils | FCGR3B,S100A9,S100A8 |
| Mast cells | TPSAB1,TPSB2,CPA3 |
| Mononuclear phagocytes | CD14,CSF1R,HLA-DRA |
| Plasmacytoid dendritic cells | IL3RA,CLEC4C,LILRA4 |

b.

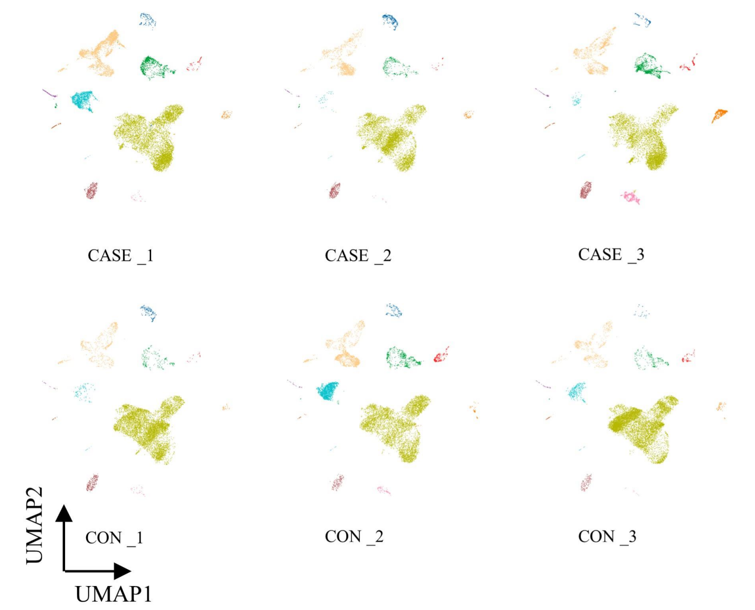

d.

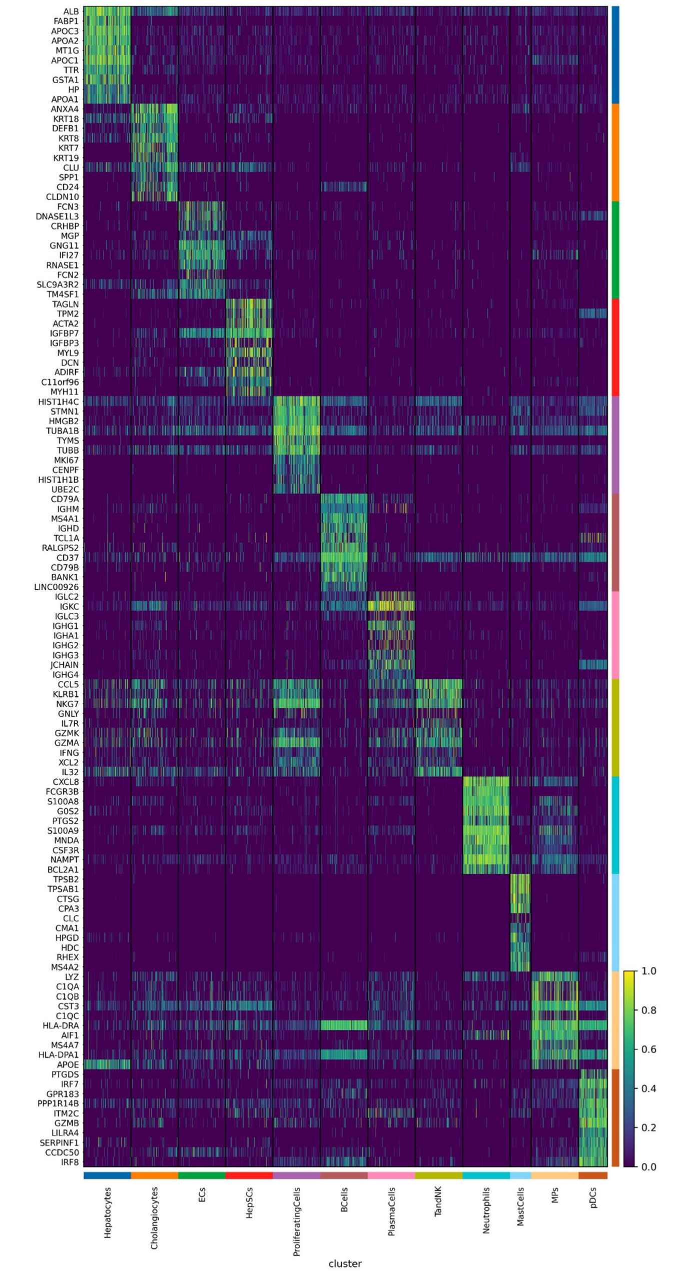

e.

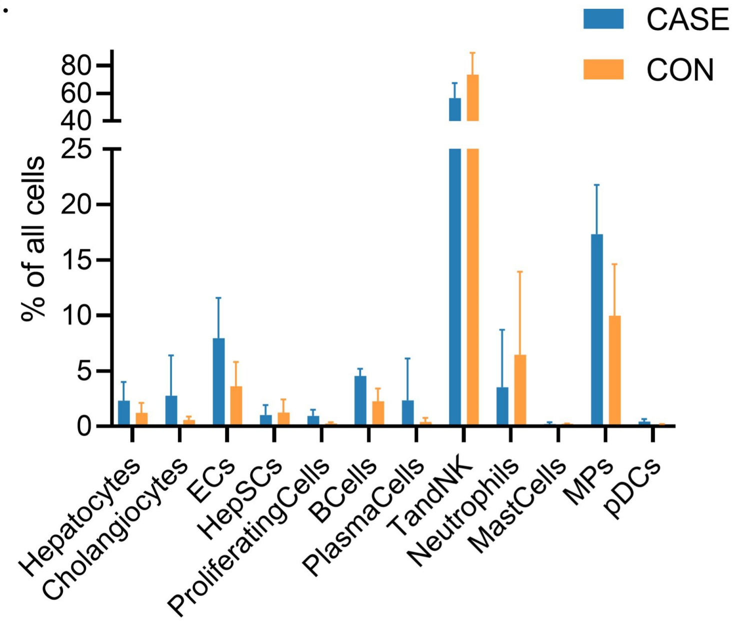

Figure S3

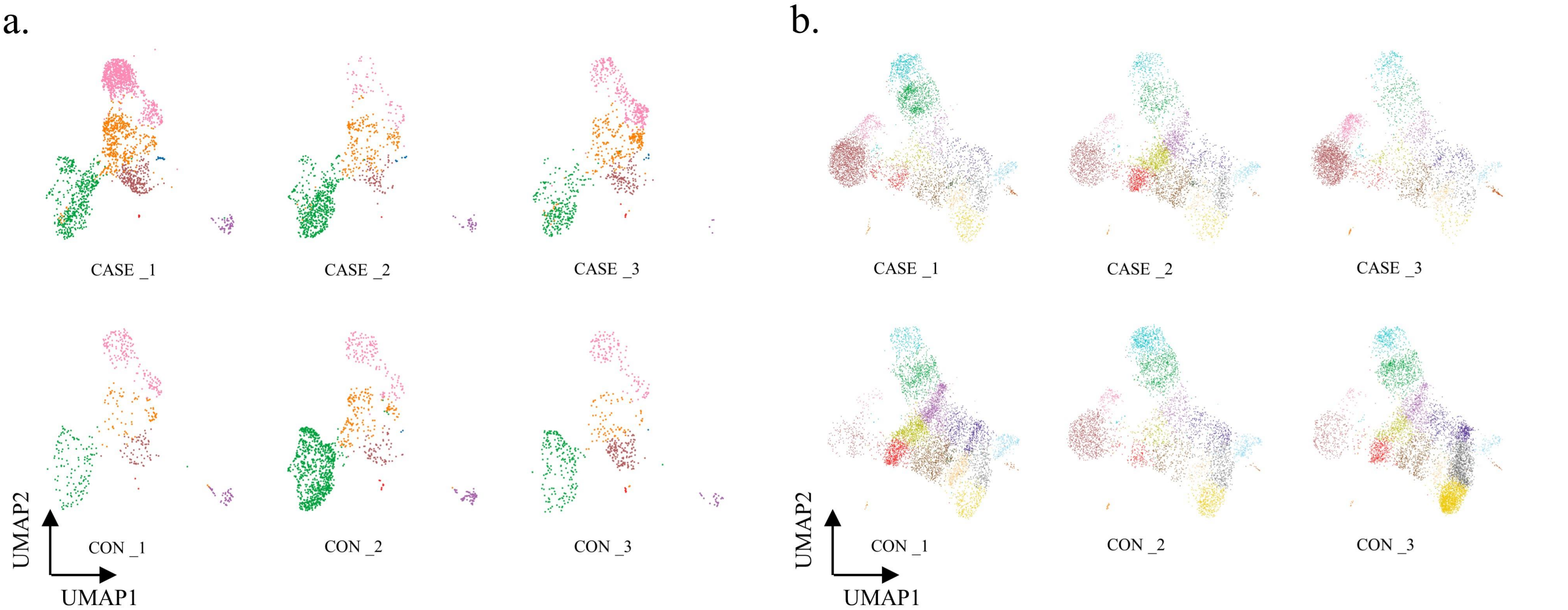

c.

**Cell marker genes for annotation of cell subtypes.**

| Celltype | Markers |
|---|---|
| Proliferating cells | MKI67,TOP2A,STMN1 |
| Macrophages | CD14,C1QA,CSF1R |
| Monocytes | LYZ,FCN1,VCAN |
| Mature dendritic cells | LAMP3,CCR7,CD83 |
| Conventional type 1 dendritic cells | CLEC9A,XCR1,IRF8 |
| Conventional type 2 dendritic cells | CD1C,CLEC10A,FCER1A |
| Kupffer cells | MARCO,CD5L,VCAM1 |
| Group 3 innate lymphoid cells | IL1R1,IL23R,KIT |
| NK T cells | NKG7,CD3D,GNLY |
| Natural killer cells | NKG7,GNLY,NCAM1 |
| CD4+ naive T cells | CCR7,SELL,LEF1 |
| CD4+ memory T cells | CD4,IL7R,CD40LG |
| CD4+ regulatory T cells | CTLA4,FOXP3,IL2RA |
| CD8+ mucosal-associated invariant T cells | KLRB1,SLC4A10,NCR3 |
| CD8+ effector T cells | CD8A,GZMK,CD3D |

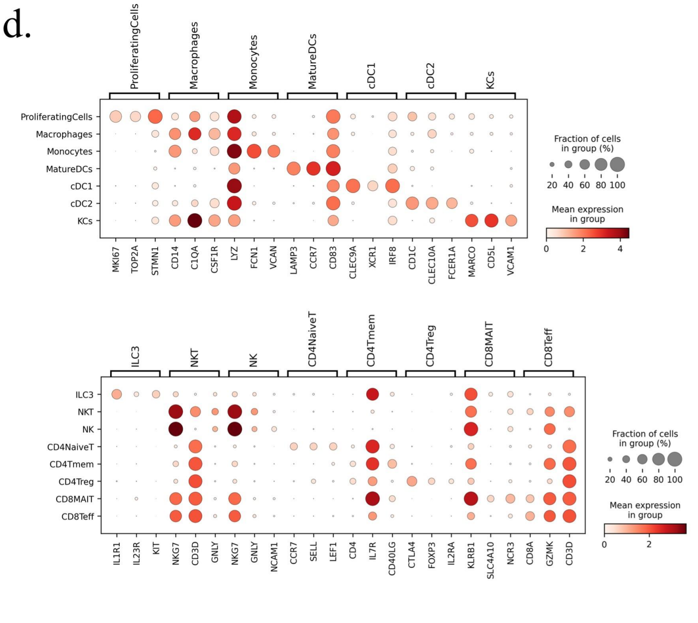

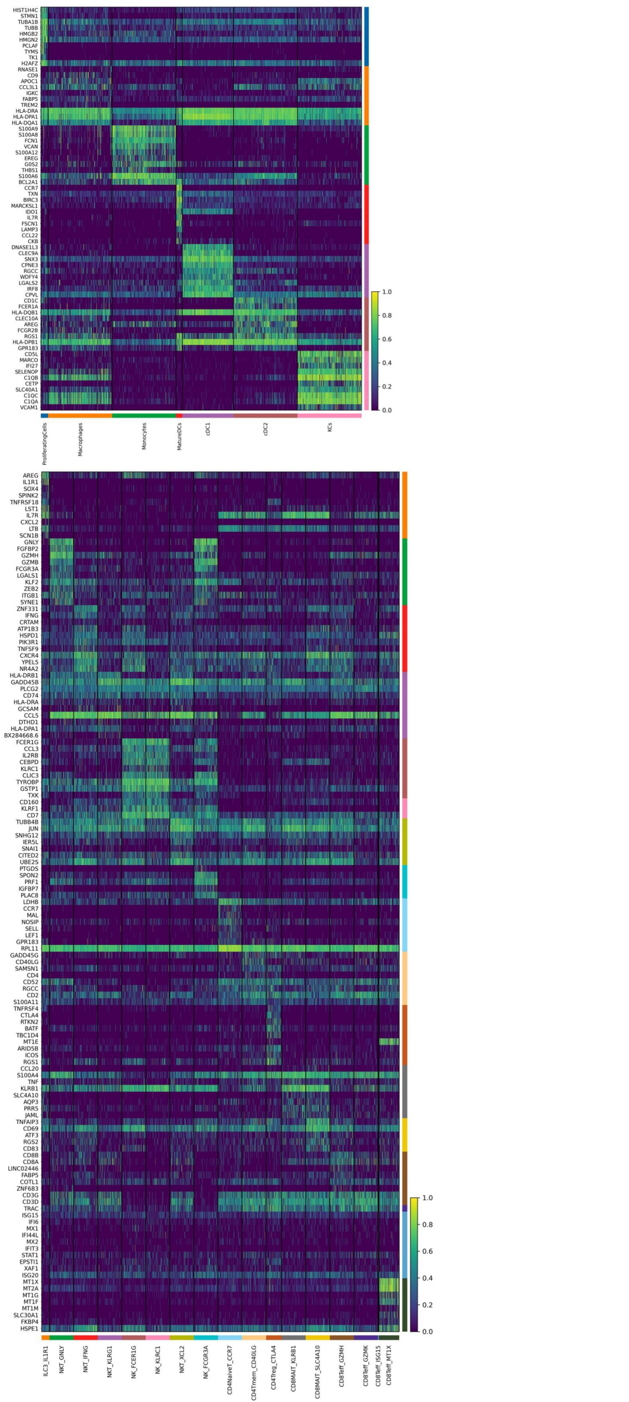

Figure S4

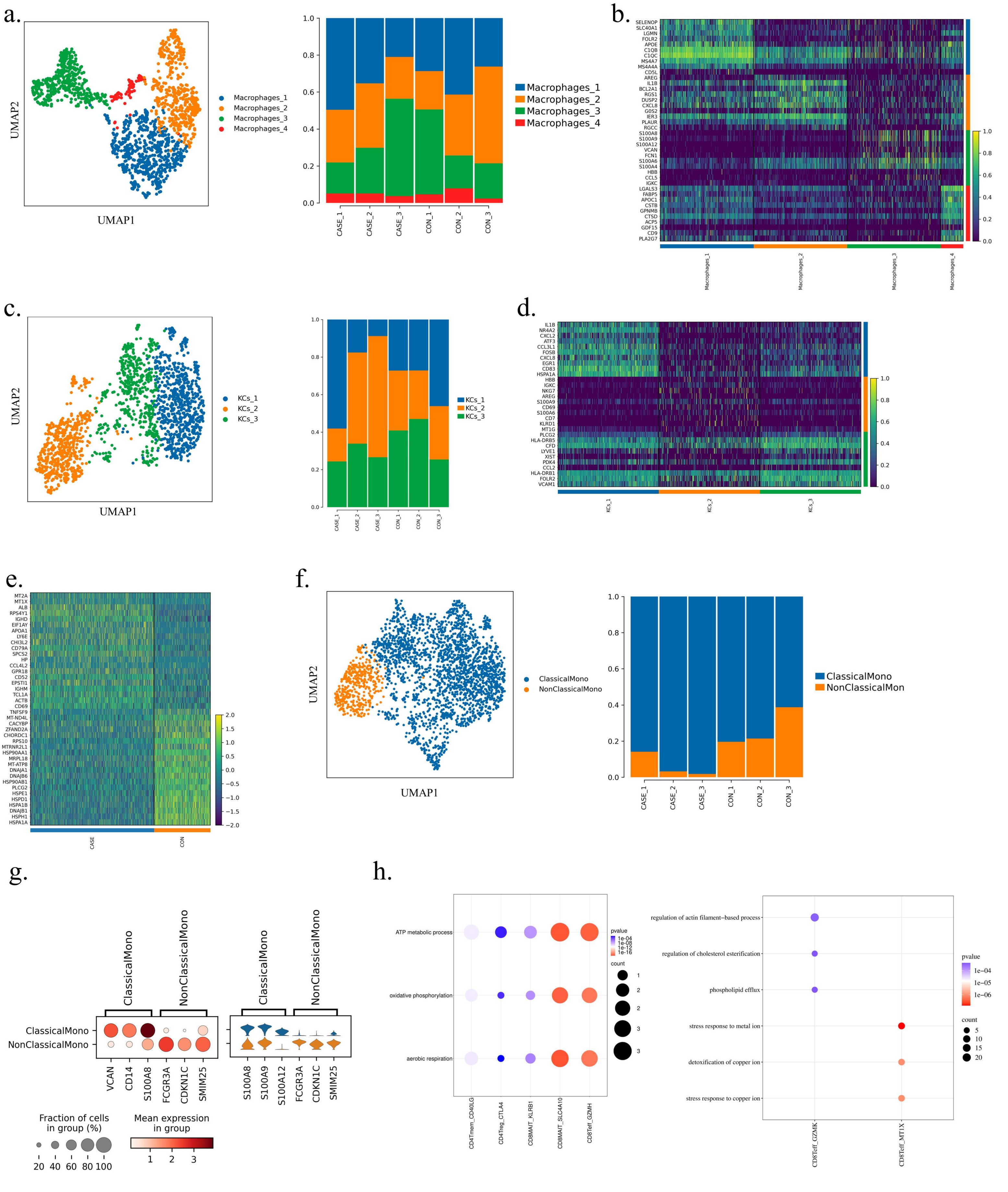
a.
UMAP2
UMAP1
Macrophages_1
Macrophages_2
Macrophages_3
Macrophages_4
CASE_1
CASE_2
CASE_3
CON_1
CON_2
CON_3
b.
c.
KCs_1
KCs_2
KCs_3
d.
e.
CASE
CON
f.
ClassicalMono
NonClassicalMono
NonClassicalMon
g.
Fraction of cells in group (%)
Mean expression in group
h.
ATP metabolic process
oxidative phosphorylation
aerobic respiration
pvalue
count
CD4Tmem_CD40LG
CD4Treg_CTLA4
CD8MAIT_KLRB1
CD8MAIT_SLC4A10
CD8Teff_GZMH
regulation of actin filament–based process
regulation of cholesterol esterification
phospholipid efflux
stress response to metal ion
detoxification of copper ion
stress response to copper ion
CD8Teff_GZMK
CD8Teff_MT1X

Figure S5

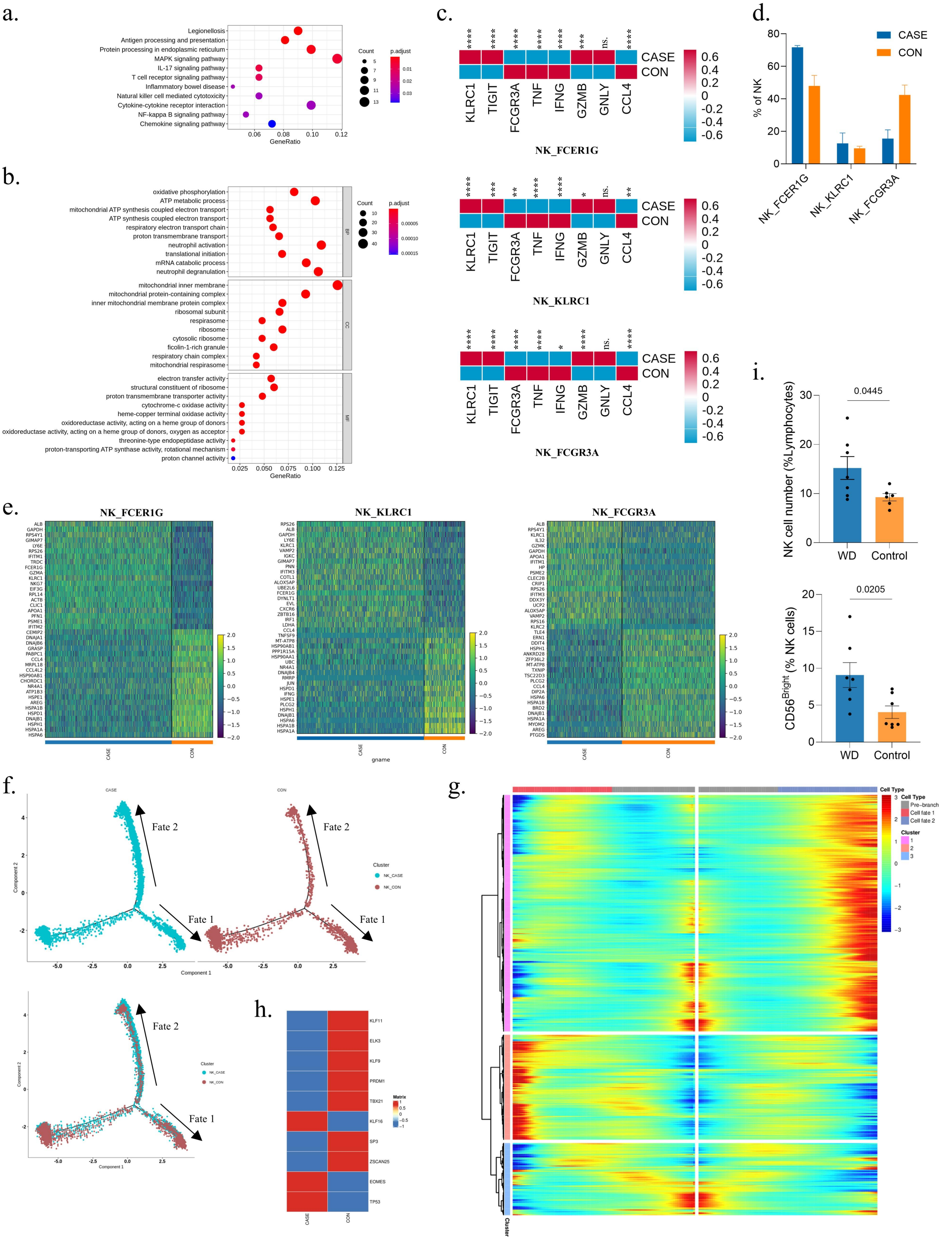
a.
Legionellosis
Antigen processing and presentation
Protein processing in endoplasmic reticulum
MAPK signaling pathway
IL-17 signaling pathway
T cell receptor signaling pathway
Inflammatory bowel disease
Natural killer cell mediated cytotoxicity
Cytokine-cytokine receptor interaction
NF-kappa B signaling pathway
Chemokine signaling pathway
GeneRatio
Count
p.adjust
b.
oxidative phosphorylation
ATP metabolic process
mitochondrial ATP synthesis coupled electron transport
ATP synthesis coupled electron transport
respiratory electron transport chain
proton transmembrane transport
neutrophil activation
translational initiation
mRNA catabolic process
neutrophil degranulation
mitochondrial inner membrane
mitochondrial protein-containing complex
inner mitochondrial membrane protein complex
ribosomal subunit
respirasome
ribosome
cytosolic ribosome
ficolin-1-rich granule
respiratory chain complex
mitochondrial respirasome
electron transfer activity
structural constituent of ribosome
proton transmembrane transporter activity
cytochrome-c oxidase activity
heme-copper terminal oxidase activity
oxidoreductase activity, acting on a heme group of donors
oxidoreductase activity, acting on a heme group of donors, oxygen as acceptor
threonine-type endopeptidase activity
proton-transporting ATP synthase activity, rotational mechanism
proton channel activity
BP
CC
MF
GeneRatio
Count
p.adjust
c.
CASE
CON
KLRC1 TIGIT FCGR3A TNF IFNG GZMB GNLY CCL4
NK_FCER1G
NK_KLRC1
NK_FCGR3A
d.
CASE
CON
% of NK
NK_FCER1G
NK_KLRC1
NK_FCGR3A
e.
NK_FCER1G
NK_KLRC1
NK_FCGR3A
CASE
CON
gname
f.
CASE
CON
Fate 1
Fate 2
Component 1
Component 2
Cluster
NK_CASE
NK_CON
g.
Cell Type
Pre-branch
Cell fate 1
Cell fate 2
Cluster
h.
KLF11
ELK3
KLF9
PRDM1
TBX21
KLF16
SP3
ZSCAN25
EOMES
TP53
Matrix
CASE
CON
i.
0.0445
NK cell number (%Lymphocytes)
WD
Control
0.0205
CD56 Bright (% NK cells)
WD
Control

Figure S6

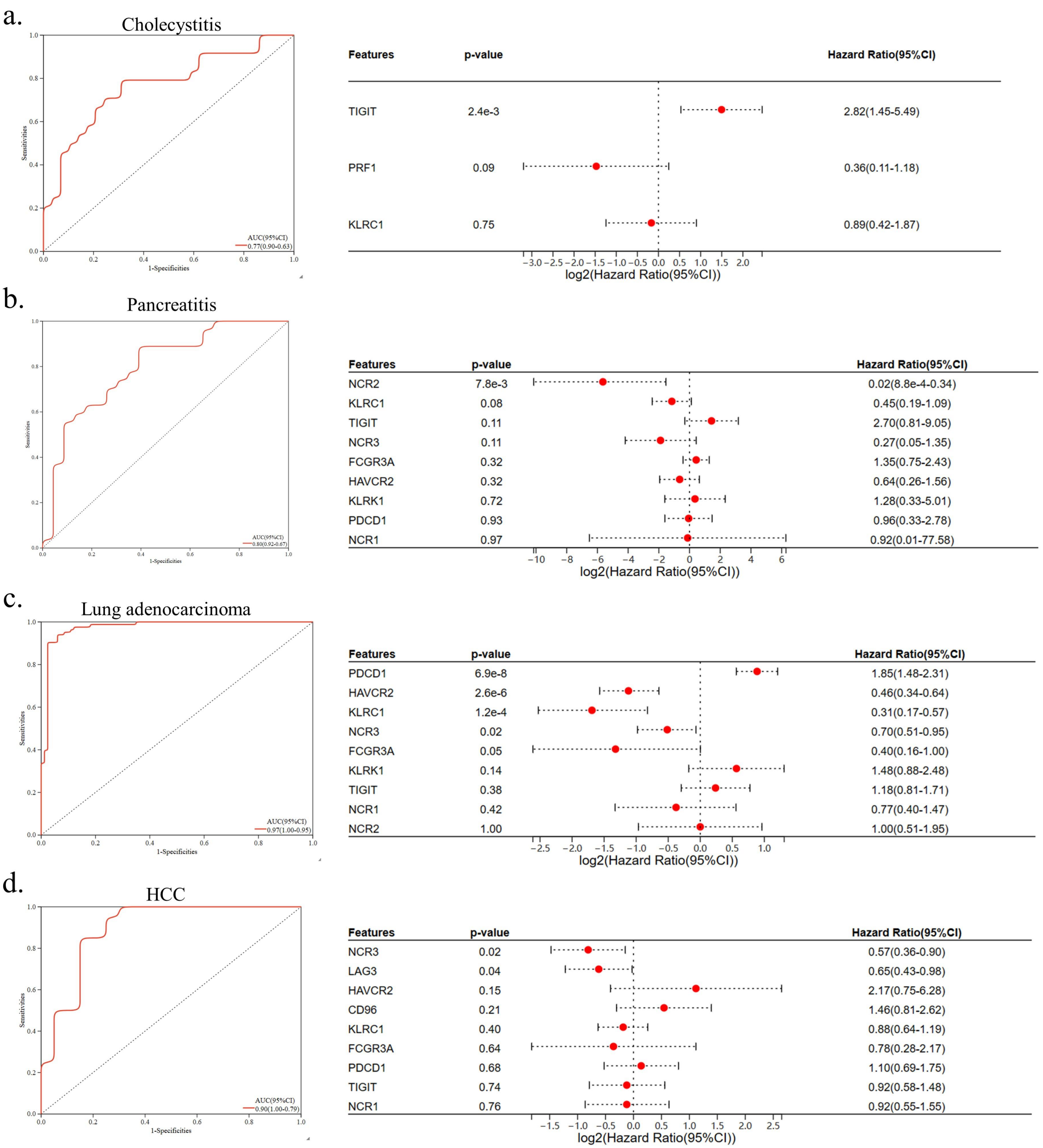
a.
Cholecystitis
Sensitivities
1-Specificities
AUC(95%CI)
0.77(0.90-0.63)
Features p-value Hazard Ratio(95%CI)
TIGIT 2.4e-3 2.82(1.45-5.49)
PRF1 0.09 0.36(0.11-1.18)
KLRC1 0.75 0.89(0.42-1.87)
log2(Hazard Ratio(95%CI))
b.
Pancreatitis
Sensitivities
1-Specificities
AUC(95%CI)
0.80(0.92-0.67)
Features p-value Hazard Ratio(95%CI)
NCR2 7.8e-3 0.02(8.8e-4-0.34)
KLRC1 0.08 0.45(0.19-1.09)
TIGIT 0.11 2.70(0.81-9.05)
NCR3 0.11 0.27(0.05-1.35)
FCGR3A 0.32 1.35(0.75-2.43)
HAVCR2 0.32 0.64(0.26-1.56)
KLRK1 0.72 1.28(0.33-5.01)
PDCD1 0.93 0.96(0.33-2.78)
NCR1 0.97 0.92(0.01-77.58)
log2(Hazard Ratio(95%CI))
c.
Lung adenocarcinoma
Sensitivities
1-Specificities
AUC(95%CI)
0.97(1.00-0.95)
Features p-value Hazard Ratio(95%CI)
PDCD1 6.9e-8 1.85(1.48-2.31)
HAVCR2 2.6e-6 0.46(0.34-0.64)
KLRC1 1.2e-4 0.31(0.17-0.57)
NCR3 0.02 0.70(0.51-0.95)
FCGR3A 0.05 0.40(0.16-1.00)
KLRK1 0.14 1.48(0.88-2.48)
TIGIT 0.38 1.18(0.81-1.71)
NCR1 0.42 0.77(0.40-1.47)
NCR2 1.00 1.00(0.51-1.95)
log2(Hazard Ratio(95%CI))
d.
HCC
Sensitivities
1-Specificities
AUC(95%CI)
0.90(1.00-0.79)
Features p-value Hazard Ratio(95%CI)
NCR3 0.02 0.57(0.36-0.90)
LAG3 0.04 0.65(0.43-0.98)
HAVCR2 0.15 2.17(0.75-6.28)
CD96 0.21 1.46(0.81-2.62)
KLRC1 0.40 0.88(0.64-1.19)
FCGR3A 0.64 0.78(0.28-2.17)
PDCD1 0.68 1.10(0.69-1.75)
TIGIT 0.74 0.92(0.58-1.48)
NCR1 0.76 0.92(0.55-1.55)
log2(Hazard Ratio(95%CI))

**Figure S1.** ***ATP7B*** **mutation causes metabolic abnormalities.** **a.** The table of clinical characteristics of patients in 3 cohorts at baseline. Data are mean± S.D. **b.** Mitochondrial stress (OCR) measured in *ATP7B*-KO and NC WRL 68 cell lines. Bar charts showing the statistical results of basal respiration and ATP production. Unpaired two-tailed t-test. **c.** Glycolysis stress (ECAR) measured in *ATP7B*-KO and NC WRL 68 cell lines. Bar charts showing the statistical results of glycolysis, and glycolysis capacity. Unpaired two-tailed t-test.

**Figure S2. The proportion and DEGs of each main cell type.** **a.** UMAP visualization of principal components by 6 samples. **b.** UMAP visualization of 12 main cell types by 6 samples, corresponding to Figure 2d. **c.** The table displaying cell marker genes for annotation of main cell types. **d.** The heat map showing the top 10 differently expressed genes (DEGs) for each cell type. **e.** The bar chart showing the average percentage of different cell types in CASE (n=3) and CON (n=3). Data are mean± S.D.

**Figure S3. The gene expression characteristics of each cell subtype.** **a.** UMAP visualization of cell subtypes in MPs by 6 samples, corresponding to Figure 3a. **b.** UMAP visualization of cell subtypes in T and NK cells by 6 samples, corresponding to Figure 3b. **c.** The table displaying cell marker genes for annotation of cell subtypes. **d.** The dot plot showing the expression of top 3 marker genes in each cell subtypes. **e.** The heatmap showing the top 10 differently expressed genes (DEGs) for each cell subtypes.

**Figure S4. The difference in percentage and gene expression of each cell subtype between CASE and CON.** **a.** UMAP visualization of the 4 further clustered subtypes of macrophages (left). The bar chart showing the percentage of each subtype across different samples (right). **b.** The heatmap showing the top 10 differently expressed genes (DEGs) for each cell subtype of macrophages. **c.** UMAP visualization of the 3 further clustered subtypes of KCs (left). The bar chart showing the percentage of each subtype across different samples (right). **d.** The heatmap showing the top 10 DEGs for each cell subtype of KCs. **e.** The heatmap showing the top 20 differently expressed genes (DEGs) for B cells. **f.** UMAP visualization of the 2 further clustered subtypes of monocytes (left). The bar chart showing the percentage of each subtype across different samples (right). **g.** The dot plot showing the expression of top 3 marker genes in each cell subtypes (left). The violin plot showing the expression of top 3 DEGs for two cell subtypes (right). **h.** The dot plot showing the upregulated biological process of GO enrichment on DEGs in each cell subtype between CASE and CON.

**Figure S5. Identification of NK cell exhaustion in WD patients.** **a.** The dot plot showing KEGG enrichment on differentially expressed genes (DEGs) in NK cells between CASE and CON. **b.** The dot plot showing GO enrichment on DEGs in NK cells between CASE and CON. BP, biological processes; CC, cellular components; MF, molecular function. **c.** The heatmap showing the relative expression level of genes in 3 NK subtypes. The color represents the relative expression level. ns., not significant; *, $P<0.05$; **, $P<0.01$; ***, $P<0.001$; ****, $P<0.0001$; Wilcox test. **d.** The bar chart showing the proportion of 3 subtypes of NK cells in the two group. **e.** The heatmap showing the top 20 differently expressed genes (DEGs) for 3 subtypes of NK cells. **f.** The plot showing trajectories of NK cells in CASE and CON. Upper: Respective trajectory. Down: Merged trajectory. **g.** The heatmap showing the expression of genes in 3 clusters in pseudotime. The cell fate is divided into two directions, and left represents fate 1 and right represents fate 2. The expression level of genes in 3 clusters was represented by colors. See full gene list in Supplementary Table S2. **h.** The heatmap showing the average expression of top 10 regulons in NK cells in CASE and CON. **i.** The bar charts showing the percentage of NK cells (WD, n=6; Control, n=7) and $CD56^{Bright}$ NK cells (WD, n=7; Control, n=7). Data are mean± S.D. Unpaired two-tailed t-test.

**Figure S6. Predictive model construction for NK cell exhaustion.** **a.-d.** Receiver operating characteristic (ROC) curve analysis of prognostic-related markers of NK cell exhaustion and corresponding AUC values for the expression cohorts (left). Forest plot of prognostic-related markers of NK cell exhaustion based on univariate Cox regression analysis (right).

**Supplementary Table S1**

**The number of each celltype**

| | **cluster** | **CASE_1** | **CASE_2** | **CASE_3** | **Sum** | **CON_1** | **CON_2** | **CON_3** | **Sum** |
|---|---|---|---|---|---|---|---|---|---|
| Main | Hepatocytes | 173 | 318 | 133 | 624 | 170 | 132 | 27 | 329 |
| | Cholangiocytes | 74 | 59 | 714 | 847 | 21 | 83 | 53 | 157 |
| | ECs | 1060 | 303 | 1155 | 2518 | 187 | 574 | 237 | 998 |
| | HepSCs | 71 | 32 | 213 | 316 | 22 | 239 | 88 | 349 |
| | ProliferatingCells | 187 | 30 | 96 | 313 | 16 | 23 | 37 | 76 |
| | BCells | 467 | 370 | 498 | 1335 | 304 | 158 | 145 | 607 |
| | PlasmaCells | 32 | 10 | 687 | 729 | 33 | 74 | 8 | 115 |
| | TandNK | 6255 | 5137 | 5130 | 16522 | 6915 | 5199 | 7915 | 20029 |
| | Neutrophils | 1175 | 35 | 63 | 1273 | 95 | 1407 | 304 | 1806 |
| | MastCells | 17 | 10 | 41 | 68 | 17 | 22 | 26 | 65 |
| | MPs | 2774 | 1122 | 1488 | 5384 | 641 | 1434 | 666 | 2741 |
| | pDCs | 76 | 12 | 51 | 139 | 11 | 12 | 21 | 44 |
| | Sum | 12361 | 7438 | 10269 | 30068 | 8432 | 9357 | 9527 | 27316 |
| MPs | ProliferatingCells | 18 | 7 | 6 | 31 | 0 | 2 | 1 | 3 |
| | Macrophages | 566 | 235 | 300 | 1101 | 87 | 179 | 84 | 350 |
| | Monocytes | 505 | 595 | 287 | 1387 | 140 | 889 | 211 | 1240 |
| | MatureDCs | 5 | 1 | 6 | 12 | 2 | 8 | 7 | 17 |
| | cDC1 | 62 | 35 | 4 | 101 | 52 | 65 | 22 | 139 |
| | cDC2 | 265 | 79 | 130 | 474 | 68 | 75 | 147 | 290 |
| | KCs | 922 | 75 | 336 | 1333 | 192 | 161 | 131 | 484 |
| | Sum | 2343 | 1027 | 1069 | 4439 | 541 | 1379 | 603 | 2523 |
| TandNK | ILC3_IL1R1 | 21 | 10 | 33 | 64 | 9 | 17 | 9 | 35 |
| | NKT_GNLY | 979 | 399 | 270 | 1648 | 725 | 674 | 831 | 2230 |
| | NKT_IFNG | 288 | 524 | 132 | 944 | 639 | 201 | 446 | 1286 |
| | NKT_KLRG1 | 168 | 483 | 204 | 855 | 950 | 245 | 591 | 1786 |
| | NK_FCER1G | 1914 | 859 | 1729 | 4502 | 210 | 887 | 369 | 1466 |
| | NK_KLRC1 | 280 | 90 | 485 | 855 | 54 | 157 | 64 | 275 |
| | NKT_XCL2 | 206 | 661 | 147 | 1014 | 799 | 205 | 465 | 1469 |
| | NK_FCGR3A | 474 | 230 | 230 | 934 | 238 | 579 | 344 | 1161 |
| | CD4NaiveT_CCR7 | 197 | 192 | 109 | 498 | 248 | 260 | 128 | 636 |
| | CD4Tmem_CD40LG | 201 | 105 | 163 | 469 | 406 | 111 | 159 | 676 |
| | CD4Treg_CTLA4 | 45 | 13 | 78 | 136 | 25 | 15 | 19 | 59 |
| | CD8MAIT_KLRB1 | 263 | 273 | 169 | 705 | 356 | 430 | 1199 | 1985 |
| | CD8MAIT_SLC4A10 | 254 | 175 | 80 | 509 | 324 | 453 | 1651 | 2428 |
| | CD8Teff_GZMH | 408 | 624 | 381 | 1413 | 835 | 294 | 554 | 1683 |
| | CD8Teff_GZMK | 221 | 200 | 247 | 668 | 669 | 357 | 659 | 1685 |
| | CD8Teff_ISG15 | 6 | 0 | 1 | 7 | 1 | 2 | 1 | 4 |
| | CD8Teff_MT1X | 74 | 79 | 29 | 182 | 53 | 7 | 12 | 72 |
| | Sum | 5999 | 4917 | 4487 | 15403 | 6541 | 4894 | 7501 | 18936 |

**The proportion of each celltype %**

| | **cluster** | **CASE_1** | **CASE_2** | **CASE_3** | **Average** | **CON_1** | **CON_2** | **CON_3** | **Average** |
|---|---|---|---|---|---|---|---|---|---|
| Main | Hepatocytes | 1.400 | 4.275 | 1.295 | 2.323 | 2.016 | 1.411 | 0.283 | 1.237 |
| | Cholangiocytes | 0.599 | 0.793 | 6.953 | 2.782 | 0.249 | 0.887 | 0.556 | 0.564 |
| | ECs | 8.575 | 4.074 | 11.247 | 7.965 | 2.218 | 6.134 | 2.488 | 3.613 |
| | HepSCs | 0.574 | 0.430 | 2.074 | 1.026 | 0.261 | 2.554 | 0.924 | 1.246 |
| | ProliferatingCells | 1.513 | 0.403 | 0.935 | 0.950 | 0.190 | 0.246 | 0.388 | 0.275 |
| | BCells | 3.778 | 4.974 | 4.850 | 4.534 | 3.605 | 1.689 | 1.522 | 2.272 |
| | PlasmaCells | 0.259 | 0.134 | 6.690 | 2.361 | 0.391 | 0.791 | 0.084 | 0.422 |
| | TandNK | 50.603 | 69.064 | 49.956 | 56.541 | 82.009 | 55.563 | 83.080 | 73.550 |
| | Neutrophils | 9.506 | 0.471 | 0.613 | 3.530 | 1.127 | 15.037 | 3.191 | 6.451 |
| | MastCells | 0.138 | 0.134 | 0.399 | 0.224 | 0.202 | 0.235 | 0.273 | 0.237 |
| | MPs | 22.442 | 15.085 | 14.490 | 17.339 | 7.602 | 15.325 | 6.991 | 9.973 |
| | pDCs | 0.615 | 0.161 | 0.497 | 0.424 | 0.130 | 0.128 | 0.220 | 0.160 |
| MPs | ProliferatingCells | 0.768 | 0.682 | 0.561 | 0.670 | 0.000 | 0.145 | 0.166 | 0.104 |
| | Macrophages | 24.157 | 22.882 | 28.064 | 25.034 | 16.081 | 12.980 | 13.930 | 14.331 |
| | Monocytes | 21.554 | 57.936 | 26.848 | 35.446 | 25.878 | 64.467 | 34.992 | 41.779 |
| | MatureDCs | 0.213 | 0.097 | 0.561 | 0.291 | 0.370 | 0.580 | 1.161 | 0.704 |
| | cDC1 | 2.646 | 3.408 | 0.374 | 2.143 | 9.612 | 4.714 | 3.648 | 5.991 |
| | cDC2 | 11.310 | 7.692 | 12.161 | 10.388 | 12.569 | 5.439 | 24.378 | 14.129 |
| | KCs | 39.351 | 7.303 | 31.431 | 26.028 | 35.490 | 11.675 | 21.725 | 22.963 |
| | ILC3_IL1R1 | 0.350 | 0.203 | 0.735 | 0.430 | 0.138 | 0.347 | 0.120 | 0.202 |
| | NKT_GNLY | 16.319 | 8.115 | 6.017 | 10.150 | 11.084 | 13.772 | 11.079 | 11.978 |
| | NKT_IFNG | 4.801 | 10.657 | 2.942 | 6.133 | 9.769 | 4.107 | 5.946 | 6.607 |
| | NKT_KLRG1 | 2.800 | 9.823 | 4.546 | 5.723 | 14.524 | 5.006 | 7.879 | 9.136 |
| | NK_FCER1G | 31.905 | 17.470 | 38.534 | 29.303 | 3.211 | 18.124 | 4.919 | 8.751 |
| | NK_KLRC1 | 4.667 | 1.830 | 10.809 | 5.769 | 0.826 | 3.208 | 0.853 | 1.629 |
| | NKT_XCL2 | 3.434 | 13.443 | 3.276 | 6.718 | 12.215 | 4.189 | 6.199 | 7.534 |

| | | | | | | | | | |
|---|---|---|---|---|---|---|---|---|---|
| | NK_FCGR3A | 7.901 | 4.678 | 5.126 | 5.902 | 3.639 | 11.831 | 4.586 | 6.685 |
| TandNK | CD4NaiveT_CCR7 | 3.284 | 3.905 | 2.429 | 3.206 | 3.791 | 5.313 | 1.706 | 3.604 |
| | CD4Tmem_CD40LG | 3.351 | 2.135 | 3.633 | 3.040 | 6.207 | 2.268 | 2.120 | 3.532 |
| | CD4Treg_CTLA4 | 0.750 | 0.264 | 1.738 | 0.918 | 0.382 | 0.306 | 0.253 | 0.314 |
| | CD8MAIT_KLRB1 | 4.384 | 5.552 | 3.766 | 4.568 | 5.443 | 8.786 | 15.985 | 10.071 |
| | CD8MAIT_SLC4A10 | 4.234 | 3.559 | 1.783 | 3.192 | 4.953 | 9.256 | 22.010 | 12.073 |
| | CD8Teff_GZMH | 6.801 | 12.691 | 8.491 | 9.328 | 12.766 | 6.007 | 7.386 | 8.720 |
| | CD8Teff_GZMK | 3.684 | 4.068 | 5.505 | 4.419 | 10.228 | 7.295 | 8.785 | 8.769 |
| | CD8Teff_ISG15 | 0.100 | 0.000 | 0.022 | 0.041 | 0.015 | 0.041 | 0.013 | 0.023 |
| | CD8Teff_MT1X | 1.234 | 1.607 | 0.646 | 1.162 | 0.810 | 0.143 | 0.160 | 0.371 |

**Supplementary Table S2**

| | gene_id | Cluster | | gene_id | Cluster | | gene_id | Cluster |
|---|---|---|---|---|---|---|---|---|
| 1 | ACTB | 1 | 757 | ARMC6 | 1 | 1517 | EGR2 | 2 |
| 2 | RPS26 | 1 | 758 | DHPS | 1 | 1518 | BTG3 | 2 |
| 3 | GIMAP7 | 1 | 759 | ELP3 | 1 | 1519 | HIC1 | 2 |
| 4 | IGKC | 1 | 760 | MRI1 | 1 | 1520 | GZF1 | 2 |
| 5 | FCER1G | 1 | 761 | CD38 | 1 | 1521 | PTGES3 | 2 |
| 6 | GSTP1 | 1 | 762 | AC022021.1 | 1 | 1522 | HMGCS1 | 2 |
| 7 | ACTG1 | 1 | 763 | PNKD | 1 | 1523 | ARG2 | 2 |
| 8 | CCL5 | 1 | 764 | C12orf10 | 1 | 1524 | SLC25A33 | 2 |
| 9 | AC007952.4 | 1 | 765 | FBXL6 | 1 | 1525 | CNOT2 | 2 |
| 10 | KLRD1 | 1 | 766 | CLUAP1 | 1 | 1526 | UBE2B | 2 |
| 11 | CLIC3 | 1 | 767 | OFD1 | 1 | 1527 | AHSA1 | 2 |
| 12 | CD7 | 1 | 768 | COX5A | 1 | 1528 | OAZ1 | 2 |
| 13 | TMSB4X | 1 | 769 | ANKRD10 | 1 | 1529 | HNRNPC | 2 |
| 14 | GZMA | 1 | 770 | ATP5S | 1 | 1530 | AMD1 | 2 |
| 15 | RSRP1 | 1 | 771 | TBC1D10A | 1 | 1531 | CHD4 | 2 |
| 16 | GAPDH | 1 | 772 | METTL26 | 1 | 1532 | MIDN | 2 |
| 17 | MYL12A | 1 | 773 | TARSL2 | 1 | 1533 | SERP1 | 2 |
| 18 | KLRC1 | 1 | 774 | TAF9 | 1 | 1534 | NAA50 | 2 |
| 19 | CD247 | 1 | 775 | ADD1 | 1 | 1535 | MOB3A | 2 |
| 20 | ID2 | 1 | 776 | DEXI | 1 | 1536 | KBTBD2 | 2 |
| 21 | XIST | 1 | 777 | SYNJ2BP | 1 | 1537 | YME1L1 | 2 |
| 22 | PFN1 | 1 | 778 | TMEM106C | 1 | 1538 | SLC2A3 | 2 |
| 23 | CLIC1 | 1 | 779 | TMOD3 | 1 | 1539 | MAT2A | 2 |
| 24 | SOCS1 | 1 | 780 | CLNS1A | 1 | 1540 | AC103591.3 | 2 |
| 25 | GZMK | 1 | 781 | STUB1 | 1 | 1541 | BIRC2 | 2 |
| 26 | CORO1A | 1 | 782 | DPM3 | 1 | 1542 | TNFSF14 | 2 |
| 27 | HCST | 1 | 783 | ATP6V1F | 1 | 1543 | POLR2K | 2 |
| 28 | HLA-B | 1 | 784 | ZNF567 | 1 | 1544 | EHD1 | 2 |
| 29 | TXK | 1 | 785 | ECHS1 | 1 | 1545 | CLK1 | 2 |
| 30 | CITED2 | 1 | 786 | NAGLU | 1 | 1546 | KPNA2 | 2 |
| 31 | ATP5F1E | 1 | 787 | MAF1 | 1 | 1547 | PABPC4 | 2 |
| 32 | TYROBP | 1 | 788 | ATP5F1C | 1 | 1548 | SUB1 | 2 |
| 33 | IFITM2 | 1 | 789 | LBR | 1 | 1549 | CCT4 | 2 |
| 34 | LY6E | 1 | 790 | TEFM | 1 | 1550 | TSC22D2 | 2 |
| 35 | SESN1 | 1 | 791 | APH1A | 1 | 1551 | RPLP1 | 2 |
| 36 | LSP1 | 1 | 792 | POLR2I | 1 | 1552 | MIR222HG | 2 |
| 37 | CD63 | 1 | 793 | CAMK1 | 1 | 1553 | SFPQ | 2 |
| 38 | CTSD | 1 | 794 | RTRAF | 1 | 1554 | RPL35 | 2 |
| 39 | EVL | 1 | 795 | BTF3L4 | 1 | 1555 | TIFA | 2 |
| 40 | PSME1 | 1 | 796 | ZFP14 | 1 | 1556 | SKP1 | 2 |
| 41 | FKBP5 | 1 | 797 | METTL23 | 1 | 1557 | TMEM263 | 2 |
| 42 | CFL1 | 1 | 798 | FAM50B | 1 | 1558 | NBEAL1 | 2 |
| 43 | TMIGD2 | 1 | 799 | COMMD9 | 1 | 1559 | UBE2A | 2 |
| 44 | GADD45G | 1 | 800 | CSK | 1 | 1560 | ZBTB21 | 2 |
| 45 | GABPB1-AS | 1 | 801 | SDAD1 | 1 | 1561 | LEPROTL1 | 2 |
| 46 | RAC2 | 1 | 802 | TINF2 | 1 | 1562 | ZBTB10 | 2 |
| 47 | ARPC3 | 1 | 803 | IMP3 | 1 | 1563 | FAM107B | 2 |
| 48 | GADD45B | 1 | 804 | SF3B5 | 1 | 1564 | TAF13 | 2 |
| 49 | ARRDC3 | 1 | 805 | APEX1 | 1 | 1565 | GOLGB1 | 2 |
| 50 | CD160 | 1 | 806 | ZNF428 | 1 | 1566 | SERTAD3 | 2 |
| 51 | ETS1 | 1 | 807 | METTL3 | 1 | 1567 | SLC1A5 | 2 |
| 52 | MYL12B | 1 | 808 | CFAP97 | 1 | 1568 | RFX1 | 2 |
| 53 | PTGDR | 1 | 809 | TERF2IP | 1 | 1569 | AZIN1 | 2 |
| 54 | AC104506.1 | 1 | 810 | STX10 | 1 | 1570 | ZDBF2 | 2 |
| 55 | ZBTB16 | 1 | 811 | RALY | 1 | 1571 | NSMCE3 | 2 |
| 56 | MAPK1 | 1 | 812 | TRAPPC2L | 1 | 1572 | MKNK2 | 2 |
| 57 | IGHA1 | 1 | 813 | CENPT | 1 | 1573 | BCL2L11 | 2 |
| 58 | RARRES3 | 1 | 814 | SLC7A6OS | 1 | 1574 | NFKB2 | 2 |
| 59 | INTS6 | 1 | 815 | C11orf58 | 1 | 1575 | H2AFJ | 2 |
| 60 | FUS | 1 | 816 | ZNF75A | 1 | 1576 | NFKBIE | 2 |
| 61 | ILF3-DT | 1 | 817 | TUBB | 1 | 1577 | UBE2D3 | 2 |
| 62 | TRAPPC1 | 1 | 818 | VPS26A | 1 | 1578 | RHOH | 2 |
| 63 | GZMM | 1 | 819 | BCL2 | 1 | 1579 | THUMPD3- | 2 |
| 64 | RASA2 | 1 | 820 | PDCD6 | 1 | 1580 | IFITM1 | 2 |
| 65 | EIF3G | 1 | 821 | SP110 | 1 | 1581 | TOB2 | 2 |
| 66 | AL355075.4 | 1 | 822 | ITGA4 | 1 | 1582 | EGR3 | 2 |

| | | | | | | | | |
|---|---|---|---|---|---|---|---|---|
| 67 | SERF2 | 1 | 823 | MLEC | 1 | 1583 | NASP | 2 |
| 68 | GIMAP4 | 1 | 824 | ASAH1 | 1 | 1584 | SIRT1 | 2 |
| 69 | ITM2C | 1 | 825 | TMEM203 | 1 | 1585 | NAMPT | 2 |
| 70 | TMA7 | 1 | 826 | C8orf59 | 1 | 1586 | RIPK2 | 2 |
| 71 | TRDC | 1 | 827 | DDIT4 | 1 | 1587 | ICAM4 | 2 |
| 72 | C4orf3 | 1 | 828 | TMEM42 | 1 | 1588 | ALG13 | 2 |
| 73 | CXCR6 | 1 | 829 | CLNK | 1 | 1589 | THAP2 | 2 |
| 74 | NDUFA4 | 1 | 830 | PGBD4 | 1 | 1590 | GCH1 | 2 |
| 75 | TLN1 | 1 | 831 | SPCS1 | 1 | 1591 | IFNGR1 | 2 |
| 76 | GIMAP1 | 1 | 832 | CDC16 | 1 | 1592 | HIST2H2AC | 2 |
| 77 | PSME2 | 1 | 833 | TSPAN5 | 1 | 1593 | PDE7A | 2 |
| 78 | UQCR11 | 1 | 834 | TRMT10B | 1 | 1594 | SRSF7 | 2 |
| 79 | PPIB | 1 | 835 | AUTS2 | 1 | 1595 | PNPLA8 | 2 |
| 80 | ELOB | 1 | 836 | TBRG4 | 1 | 1596 | MMP9 | 2 |
| 81 | GGNBP2 | 1 | 837 | PLP2 | 1 | 1597 | RBM38 | 2 |
| 82 | SPCS2 | 1 | 838 | XRCC5 | 1 | 1598 | ATG101 | 2 |
| 83 | CHST12 | 1 | 839 | AC243829.1 | 1 | 1599 | NLRP3 | 2 |
| 84 | RHOC | 1 | 840 | NIFK | 1 | 1600 | CASP3 | 2 |
| 85 | PRELID1 | 1 | 841 | ZNF302 | 1 | 1601 | JPT1 | 2 |
| 86 | ARHGDIB | 1 | 842 | ARSK | 1 | 1602 | TANK | 2 |
| 87 | AP001160.1 | 1 | 843 | SNAPC5 | 1 | 1603 | ISG20L2 | 2 |
| 88 | AL627171.1 | 1 | 844 | GHITM | 1 | 1604 | CNOT6L | 2 |
| 89 | RHOA | 1 | 845 | COMMD2 | 1 | 1605 | SIK1B | 2 |
| 90 | ENO1 | 1 | 846 | COMTD1 | 1 | 1606 | RHEB | 2 |
| 91 | IKZF1 | 1 | 847 | WDR1 | 1 | 1607 | LIF | 2 |
| 92 | AC108134.2 | 1 | 848 | DNMT3A | 1 | 1608 | SC5D | 2 |
| 93 | PPP1CA | 1 | 849 | SYK | 1 | 1609 | MAP1LC3A | 2 |
| 94 | PPP1R14B | 1 | 850 | TMEM126B | 1 | 1610 | USP36 | 2 |
| 95 | ATP5MC2 | 1 | 851 | U2AF1L5 | 1 | 1611 | RAB21 | 2 |
| 96 | IGHM | 1 | 852 | TMCO1 | 1 | 1612 | RYBP | 2 |
| 97 | CDC42 | 1 | 853 | MPC1 | 1 | 1613 | TFRC | 2 |
| 98 | ACAP1 | 1 | 854 | CIR1 | 1 | 1614 | DDX21 | 2 |
| 99 | IFI16 | 1 | 855 | RBL2 | 1 | 1615 | ELOVL5 | 2 |
| 100 | OST4 | 1 | 856 | CCDC124 | 1 | 1616 | DBF4 | 2 |
| 101 | PNISR | 1 | 857 | SFXN1 | 1 | 1617 | KDM2A | 2 |
| 102 | PLEKHF1 | 1 | 858 | TSNAX | 1 | 1618 | RNF168 | 2 |
| 103 | DAD1 | 1 | 859 | M6PR | 1 | 1619 | DUSP6 | 2 |
| 104 | PYHIN1 | 1 | 860 | MRPS5 | 1 | 1620 | SIK2 | 2 |
| 105 | PRDX5 | 1 | 861 | DCP2 | 1 | 1621 | DNAJB9 | 2 |
| 106 | CAP1 | 1 | 862 | CCDC85B | 1 | 1622 | AKIRIN1 | 2 |
| 107 | ALOX5AP | 1 | 863 | SH2D1B | 1 | 1623 | RARA | 2 |
| 108 | APMAP | 1 | 864 | RCBTB2 | 1 | 1624 | LRIF1 | 2 |
| 109 | BST2 | 1 | 865 | H6PD | 1 | 1625 | MAP2K3 | 2 |
| 110 | EOMES | 1 | 866 | RBM43 | 1 | 1626 | EIF1AY | 2 |
| 111 | SET | 1 | 867 | SUMO2 | 1 | 1627 | PPP1CB | 2 |
| 112 | LAT2 | 1 | 868 | OGT | 1 | 1628 | CRYBG1 | 2 |
| 113 | ARGLU1 | 1 | 869 | PSMF1 | 1 | 1629 | RASL11A | 2 |
| 114 | FASLG | 1 | 870 | ZNF439 | 1 | 1630 | MOB4 | 2 |
| 115 | CD53 | 1 | 871 | VDAC1 | 1 | 1631 | DYNLL2 | 2 |
| 116 | TBC1D10C | 1 | 872 | PARK7 | 1 | 1632 | CYSLTR2 | 2 |
| 117 | IFIT2 | 1 | 873 | A1BG | 1 | 1633 | CCNT1 | 2 |
| 118 | MSN | 1 | 874 | EMC1 | 1 | 1634 | ISCA1 | 2 |
| 119 | TMEM141 | 1 | 875 | CKLF | 1 | 1635 | CMC1 | 2 |
| 120 | ARL6IP4 | 1 | 876 | PAK2 | 1 | 1636 | SRSF5 | 2 |
| 121 | EIF5A | 1 | 877 | FNTA | 1 | 1637 | SNHG8 | 2 |
| 122 | HMGB1 | 1 | 878 | UBE2D2 | 1 | 1638 | TRA2A | 2 |
| 123 | PSMB8 | 1 | 879 | PIGBOS1 | 1 | 1639 | LUZP1 | 2 |
| 124 | PTPRC | 1 | 880 | NDUFAB1 | 1 | 1640 | NKRF | 2 |
| 125 | IGHG3 | 1 | 881 | CHRNB1 | 1 | 1641 | PPP2CA | 2 |
| 126 | C19orf66 | 1 | 882 | COX14 | 1 | 1642 | PIM2 | 2 |
| 127 | PIK3IP1 | 1 | 883 | AL451085.1 | 1 | 1643 | ISG15 | 2 |
| 128 | HMGN1 | 1 | 884 | CTBP1 | 1 | 1644 | EGR4 | 2 |
| 129 | ANXA6 | 1 | 885 | BUB3 | 1 | 1645 | MORF4L2 | 2 |
| 130 | ECH1 | 1 | 886 | NUTF2 | 1 | 1646 | NDUFS5 | 2 |
| 131 | HSD17B10 | 1 | 887 | HINT2 | 1 | 1647 | TOPORS | 2 |
| 132 | CSKMT | 1 | 888 | MRPL28 | 1 | 1648 | AL359915.2 | 2 |
| 133 | PAXX | 1 | 889 | RTP4 | 1 | 1649 | ZNF571 | 2 |
| 134 | ATP5MD | 1 | 890 | ENPP4 | 1 | 1650 | AURKA | 2 |

| | | |
|---|---|---|
| 135 | VAMP8 | 1 |
| 136 | GMFG | 1 |
| 137 | WDR83OS | 1 |
| 138 | GTF3A | 1 |
| 139 | MYL6 | 1 |
| 140 | ITGB2 | 1 |
| 141 | TRAF3IP3 | 1 |
| 142 | PSMB3 | 1 |
| 143 | COX6B1 | 1 |
| 144 | NT5C | 1 |
| 145 | GPATCH8 | 1 |
| 146 | ARPC4 | 1 |
| 147 | IGLC2 | 1 |
| 148 | CAPZB | 1 |
| 149 | MDH2 | 1 |
| 150 | COX6C | 1 |
| 151 | AC243965.1 | 1 |
| 152 | TCIRG1 | 1 |
| 153 | FKBP8 | 1 |
| 154 | PHYKPL | 1 |
| 155 | 1-Sep | 1 |
| 156 | NDUFB2 | 1 |
| 157 | LCP1 | 1 |
| 158 | CD81 | 1 |
| 159 | APBB1IP | 1 |
| 160 | COTL1 | 1 |
| 161 | AC243960.1 | 1 |
| 162 | HLA-DQB1 | 1 |
| 163 | S100A11 | 1 |
| 164 | SYNGR1 | 1 |
| 165 | SSBP4 | 1 |
| 166 | UQCR10 | 1 |
| 167 | ITM2A | 1 |
| 168 | COG2 | 1 |
| 169 | MRPS34 | 1 |
| 170 | MATK | 1 |
| 171 | IFITM3 | 1 |
| 172 | PPP1R18 | 1 |
| 173 | UBE2L6 | 1 |
| 174 | APRT | 1 |
| 175 | GSTK1 | 1 |
| 176 | TESC | 1 |
| 177 | YWHAB | 1 |
| 178 | SLFN5 | 1 |
| 179 | RASA1 | 1 |
| 180 | NEDD8 | 1 |
| 181 | DYNLT1 | 1 |
| 182 | AP3S1 | 1 |
| 183 | PARP8 | 1 |
| 184 | AC023157.3 | 1 |
| 185 | ATP5F1A | 1 |
| 186 | ARPC5 | 1 |
| 187 | ATP5F1D | 1 |
| 188 | RNF187 | 1 |
| 189 | LAMTOR4 | 1 |
| 190 | COPE | 1 |
| 191 | PLEKHJ1 | 1 |
| 192 | TMED9 | 1 |
| 193 | EID1 | 1 |
| 194 | NDUFC2 | 1 |
| 195 | ANP32A | 1 |
| 196 | PSMB1 | 1 |
| 197 | PSMB9 | 1 |
| 198 | ATP5MC3 | 1 |
| 199 | ATP5ME | 1 |
| 200 | MMP25-AS | 1 |
| 201 | FBXW5 | 1 |
| 202 | IPCEF1 | 1 |
| 891 | FARSA | 1 |
| 892 | AC017083.1 | 1 |
| 893 | MLH3 | 1 |
| 894 | NIFK-AS1 | 1 |
| 895 | BOD1 | 1 |
| 896 | SASH3 | 1 |
| 897 | ZDHHC24 | 1 |
| 898 | C19orf24 | 1 |
| 899 | CNDP2 | 1 |
| 900 | NOL7 | 1 |
| 901 | CCDC28A | 1 |
| 902 | ARHGDIA | 1 |
| 903 | ARL6IP5 | 1 |
| 904 | IER3IP1 | 1 |
| 905 | IFI35 | 1 |
| 906 | LYPLA2 | 1 |
| 907 | POLR2L | 1 |
| 908 | TMEM156 | 1 |
| 909 | EIF3K | 1 |
| 910 | HAGHL | 1 |
| 911 | NCOR1 | 1 |
| 912 | BRI3 | 1 |
| 913 | JMJD8 | 1 |
| 914 | TPR | 1 |
| 915 | VEGFB | 1 |
| 916 | OSTF1 | 1 |
| 917 | WDR77 | 1 |
| 918 | SH3BGRL | 1 |
| 919 | UTP25 | 1 |
| 920 | ORMDL2 | 1 |
| 921 | PKN1 | 1 |
| 922 | PEPD | 1 |
| 923 | BABAM1 | 1 |
| 924 | DNAJC1 | 1 |
| 925 | G6PD | 1 |
| 926 | SDHAF2 | 1 |
| 927 | NUDCD2 | 1 |
| 928 | PARP16 | 1 |
| 929 | SRP72 | 1 |
| 930 | LZTR1 | 1 |
| 931 | TMA16 | 1 |
| 932 | GNG5 | 1 |
| 933 | NAPEPLD | 1 |
| 934 | C19orf53 | 1 |
| 935 | MTERF4 | 1 |
| 936 | CUTA | 1 |
| 937 | VSTM4 | 1 |
| 938 | NAXE | 1 |
| 939 | AC110769.2 | 1 |
| 940 | MIF4GD | 1 |
| 941 | MZT2A | 1 |
| 942 | PDLIM1 | 1 |
| 943 | NELFB | 1 |
| 944 | CUEDC2 | 1 |
| 945 | EXOSC7 | 1 |
| 946 | FGD4 | 1 |
| 947 | SNRPD3 | 1 |
| 948 | POLR2J | 1 |
| 949 | STK17A | 1 |
| 950 | MESD | 1 |
| 951 | RASAL3 | 1 |
| 952 | MEA1 | 1 |
| 953 | DYNC1I2 | 1 |
| 954 | VPS36 | 1 |
| 955 | ABHD14B | 1 |
| 956 | MRPL44 | 1 |
| 957 | RNF167 | 1 |
| 958 | CCDC127 | 1 |
| 1651 | GADD45A | 2 |
| 1652 | CCDC117 | 2 |
| 1653 | TRIM39 | 2 |
| 1654 | TGIF2 | 2 |
| 1655 | ATP1A1 | 2 |
| 1656 | IRF2BP2 | 2 |
| 1657 | TMEM217 | 2 |
| 1658 | EMD | 2 |
| 1659 | ZFAS1 | 2 |
| 1660 | SLC35A2 | 2 |
| 1661 | CD72 | 2 |
| 1662 | SRSF3 | 2 |
| 1663 | BTBD7 | 2 |
| 1664 | RABGEF1 | 2 |
| 1665 | ELMSAN1 | 2 |
| 1666 | ZNF570 | 2 |
| 1667 | FOSL1 | 2 |
| 1668 | EIF4G2 | 2 |
| 1669 | DCUN1D3 | 2 |
| 1670 | MEF2D | 2 |
| 1671 | RSL24D1 | 2 |
| 1672 | TRAF4 | 2 |
| 1673 | MRPL1 | 2 |
| 1674 | CBX4 | 2 |
| 1675 | FABP5 | 2 |
| 1676 | RPL22L1 | 2 |
| 1677 | NSUN6 | 2 |
| 1678 | STIP1 | 2 |
| 1679 | SLC35D1 | 2 |
| 1680 | GLS | 2 |
| 1681 | SNU13 | 2 |
| 1682 | NDEL1 | 2 |
| 1683 | WDR47 | 2 |
| 1684 | SPATA2L | 2 |
| 1685 | AC058791.1 | 2 |
| 1686 | TARS | 2 |
| 1687 | NUDT4 | 2 |
| 1688 | SERPINH1 | 2 |
| 1689 | LMNB1 | 2 |
| 1690 | SS18L2 | 2 |
| 1691 | IL23R | 2 |
| 1692 | UBR1 | 2 |
| 1693 | SLC25A3 | 2 |
| 1694 | CD27 | 2 |
| 1695 | TMEM88 | 2 |
| 1696 | DNTTIP2 | 2 |
| 1697 | STMN1 | 2 |
| 1698 | LETM2 | 2 |
| 1699 | MASTL | 2 |
| 1700 | UBE2D1 | 2 |
| 1701 | CHIC2 | 2 |
| 1702 | AL021707.6 | 2 |
| 1703 | MAZ | 2 |
| 1704 | TOP1 | 2 |
| 1705 | SPRY1 | 2 |
| 1706 | VEGFA | 2 |
| 1707 | TP53INP2 | 2 |
| 1708 | CYLD | 2 |
| 1709 | FAM222A | 2 |
| 1710 | PDCL3 | 2 |
| 1711 | INPP4B | 2 |
| 1712 | TESK1 | 2 |
| 1713 | ALB | 2 |
| 1714 | STARD7 | 2 |
| 1715 | RNF138 | 2 |
| 1716 | PER2 | 2 |
| 1717 | AHI1 | 2 |
| 1718 | AC009404.1 | 2 |

| | | | | | | | | |
|---|---|---|---|---|---|---|---|---|
| 203 | TGFBR2 | 1 | 959 | PRR13 | 1 | 1719 | SNHG15 | 2 |
| 204 | PEBP1 | 1 | 960 | PSIP1 | 1 | 1720 | UBAP1 | 2 |
| 205 | PKM | 1 | 961 | SH3GLB2 | 1 | 1721 | INTS6L | 2 |
| 206 | TPI1 | 1 | 962 | TMEM258 | 1 | 1722 | CYGB | 2 |
| 207 | IGHG1 | 1 | 963 | MGAT4B | 1 | 1723 | CH25H | 2 |
| 208 | SNTB2 | 1 | 964 | SUMF2 | 1 | 1724 | B3GALNT2 | 2 |
| 209 | LAMTOR1 | 1 | 965 | SDHAF1 | 1 | 1725 | ADRA2B | 2 |
| 210 | LINC00324 | 1 | 966 | SRSF10 | 1 | 1726 | PHLDB2 | 2 |
| 211 | MIEN1 | 1 | 967 | HEXDC | 1 | 1727 | SEC24A | 2 |
| 212 | LAPTM5 | 1 | 968 | TAF15 | 1 | 1728 | CDC37 | 2 |
| 213 | CALM3 | 1 | 969 | OTUD6B-A | 1 | 1729 | CTDP1 | 2 |
| 214 | CCT7 | 1 | 970 | VKORC1 | 1 | 1730 | CCNYL1 | 2 |
| 215 | NME1 | 1 | 971 | C14orf119 | 1 | 1731 | ATF4 | 2 |
| 216 | RFLNB | 1 | 972 | FO393401.1 | 1 | 1732 | ARL4D | 2 |
| 217 | COX8A | 1 | 973 | LINC02084 | 1 | 1733 | DUSP10 | 2 |
| 218 | C19orf70 | 1 | 974 | TMEM134 | 1 | 1734 | KCNQ1OT1 | 2 |
| 219 | RSBN1L | 1 | 975 | TBCE | 1 | 1735 | KLF16 | 2 |
| 220 | DERL1 | 1 | 976 | DCTPP1 | 1 | 1736 | ARHGEF39 | 2 |
| 221 | POLR2G | 1 | 977 | TFB1M | 1 | 1737 | SPAG4 | 2 |
| 222 | ATP5MPL | 1 | 978 | EPS8L2 | 1 | 1738 | EZH2 | 2 |
| 223 | BCAP31 | 1 | 979 | EMC6 | 1 | 1739 | STAG2 | 2 |
| 224 | TGOLN2 | 1 | 980 | AC026471.1 | 1 | 1740 | F12 | 2 |
| 225 | UBL5 | 1 | 981 | PPIE | 1 | 1741 | MED13 | 2 |
| 226 | HBB | 1 | 982 | CRAMP1 | 1 | 1742 | RAB11FIP1 | 2 |
| 227 | ATP5MF | 1 | 983 | THUMPD3 | 1 | 1743 | EML4 | 2 |
| 228 | RNF213 | 1 | 984 | AKT1 | 1 | 1744 | RNMT | 2 |
| 229 | CEBPD | 1 | 985 | MIGA1 | 1 | 1745 | IRGQ | 2 |
| 230 | HMOX2 | 1 | 986 | SEC22B | 1 | 1746 | GEM | 2 |
| 231 | LINC01970 | 1 | 987 | DENND6A- | 1 | 1747 | TENT4B | 2 |
| 232 | CHMP2A | 1 | 988 | ADGRG3 | 1 | 1748 | WDR26 | 2 |
| 233 | COPS9 | 1 | 989 | LPIN2 | 1 | 1749 | TGIF1 | 2 |
| 234 | NUCKS1 | 1 | 990 | NAP1L4 | 1 | 1750 | WDR45B | 2 |
| 235 | ACTR2 | 1 | 991 | ZNF326 | 1 | 1751 | NFAT5 | 2 |
| 236 | FERMT3 | 1 | 992 | AL358472.4 | 1 | 1752 | IL27RA | 2 |
| 237 | FIS1 | 1 | 993 | FANCF | 1 | 1753 | HELB | 2 |
| 238 | NABP1 | 1 | 994 | MAPKAPK5 | 1 | 1754 | ARID4B | 2 |
| 239 | CSTB | 1 | 995 | STXBP2 | 1 | 1755 | SKI | 2 |
| 240 | TRBC2 | 1 | 996 | HIGD1A | 1 | 1756 | CGAS | 2 |
| 241 | 6-Sep | 1 | 997 | GYPC | 1 | 1757 | RNF125 | 2 |
| 242 | HIST1H2BN | 1 | 998 | COA1 | 1 | 1758 | BRMS1L | 2 |
| 243 | ADH5 | 1 | 999 | TMEM179B | 1 | 1759 | EIF3J | 2 |
| 244 | PSMB10 | 1 | 1000 | VPS13C | 1 | 1760 | CCL20 | 2 |
| 245 | H1FX | 1 | 1001 | FEZ2 | 1 | 1761 | HBP1 | 2 |
| 246 | AP2M1 | 1 | 1002 | RNPC3 | 1 | 1762 | BRD1 | 2 |
| 247 | MCTP2 | 1 | 1003 | SRSF8 | 1 | 1763 | RPL8 | 2 |
| 248 | ACTR3 | 1 | 1004 | JCHAIN | 1 | 1764 | PNP | 2 |
| 249 | SDF2L1 | 1 | 1005 | KLRF1 | 1 | 1765 | NRBF2 | 2 |
| 250 | COQ7 | 1 | 1006 | TMEM256 | 1 | 1766 | SESN2 | 2 |
| 251 | ATP5PO | 1 | 1007 | RBM25 | 1 | 1767 | CXCL2 | 2 |
| 252 | ETFB | 1 | 1008 | FYN | 1 | 1768 | CYB5D1 | 2 |
| 253 | TSTD1 | 1 | 1009 | ARL16 | 1 | 1769 | WHRN | 2 |
| 254 | ATP6V0E1 | 1 | 1010 | DRG2 | 1 | 1770 | TRA2B | 2 |
| 255 | AL645728.1 | 1 | 1011 | LUC7L3 | 1 | 1771 | STAT4 | 2 |
| 256 | DBNL | 1 | 1012 | ANGEL2 | 1 | 1772 | S1PR2 | 2 |
| 257 | DENND2D | 1 | 1013 | AP1M1 | 1 | 1773 | RBBP6 | 2 |
| 258 | TMEM109 | 1 | 1014 | SPSB3 | 1 | 1774 | MAGOH | 2 |
| 259 | NDUFA3 | 1 | 1015 | CCDC115 | 1 | 1775 | RHOB | 2 |
| 260 | IGFBP2 | 1 | 1016 | PLD3 | 1 | 1776 | STX17-AS1 | 2 |
| 261 | GNAI2 | 1 | 1017 | AC099778.1 | 1 | 1777 | N4BP2L1 | 2 |
| 262 | AL135925.1 | 1 | 1018 | AC012360.3 | 1 | 1778 | PLEKHA2 | 2 |
| 263 | SRM | 1 | 1019 | SEM1 | 1 | 1779 | LTA | 2 |
| 264 | TMEM230 | 1 | 1020 | VTI1B | 1 | 1780 | LYSMD3 | 2 |
| 265 | APOBEC3G | 1 | 1021 | THOC7 | 1 | 1781 | IL1R2 | 2 |
| 266 | HNRNPD | 1 | 1022 | AK9 | 1 | 1782 | MRFAP1 | 2 |
| 267 | MZT2B | 1 | 1023 | PPCS | 1 | 1783 | HIST1H1A | 2 |
| 268 | HCLS1 | 1 | 1024 | MRPS33 | 1 | 1784 | LINC00513 | 2 |
| 269 | PPP1R2C | 1 | 1025 | PCYT2 | 1 | 1785 | KANSL2 | 2 |
| 270 | BSG | 1 | 1026 | ATP5PB | 1 | 1786 | AC144831.1 | 2 |

| | | | | | | | | |
|---|---|---|---|---|---|---|---|---|
| 271 | TWF2 | 1 | 1027 | TMX3 | 1 | 1787 | DRAM1 | 2 |
| 272 | RBM3 | 1 | 1028 | MBD2 | 1 | 1788 | RBM12 | 2 |
| 273 | TMED4 | 1 | 1029 | PPP1R3E | 1 | 1789 | SNAPC1 | 2 |
| 274 | ABI3 | 1 | 1030 | PDZD11 | 1 | 1790 | BEX5 | 2 |
| 275 | BRK1 | 1 | 1031 | SLC35B1 | 1 | 1791 | CXorf40A | 2 |
| 276 | SMC1A | 1 | 1032 | PMS1 | 1 | 1792 | TXLNG | 2 |
| 277 | PSMA5 | 1 | 1033 | RNASEH2C | 1 | 1793 | G0S2 | 2 |
| 278 | DDX46 | 1 | 1034 | EIF4A2 | 1 | 1794 | CD44 | 2 |
| 279 | CD74 | 1 | 1035 | TMEM14C | 1 | 1795 | NARF | 2 |
| 280 | PYCARD | 1 | 1036 | SPATC1L | 1 | 1796 | C2orf40 | 2 |
| 281 | PHPT1 | 1 | 1037 | PEX13 | 1 | 1797 | PPP2R2A | 2 |
| 282 | MINOS1 | 1 | 1038 | HIPK3 | 1 | 1798 | NSD3 | 2 |
| 283 | CFLAR | 1 | 1039 | AAK1 | 1 | 1799 | ALAS1 | 2 |
| 284 | PIP4K2A | 1 | 1040 | BOD1L1 | 1 | 1800 | SPAG1 | 2 |
| 285 | PREX1 | 1 | 1041 | CBWD3 | 1 | 1801 | DDX18 | 2 |
| 286 | PRTN3 | 1 | 1042 | THEMIS2 | 1 | 1802 | INSIG1 | 2 |
| 287 | SP100 | 1 | 1043 | AL391121.1 | 1 | 1803 | MYLIP | 2 |
| 288 | ATP5PF | 1 | 1044 | CABIN1 | 1 | 1804 | HES1 | 2 |
| 289 | ATP5MC1 | 1 | 1045 | SAP18 | 1 | 1805 | PAF1 | 2 |
| 290 | IL18 | 1 | 1046 | TLK1 | 1 | 1806 | CAAP1 | 2 |
| 291 | AC012306.2 | 1 | 1047 | SMG6 | 1 | 1807 | RGS13 | 2 |
| 292 | IKZF3 | 1 | 1048 | STOM | 1 | 1808 | AC016831.1 | 2 |
| 293 | C9orf16 | 1 | 1049 | PPP1CC | 1 | 1809 | COX7B | 2 |
| 294 | MAP4 | 1 | 1050 | MRPS12 | 1 | 1810 | CRY1 | 2 |
| 295 | SNX17 | 1 | 1051 | VAMP4 | 1 | 1811 | BUD31 | 2 |
| 296 | IL2RB | 1 | 1052 | H2AFY | 1 | 1812 | PTS | 2 |
| 297 | RPA2 | 1 | 1053 | LYRM2 | 1 | 1813 | RNF19B | 2 |
| 298 | VAMP2 | 1 | 1054 | HACL1 | 1 | 1814 | AL021707.1 | 2 |
| 299 | AL021453.1 | 1 | 1055 | DDX39B | 1 | 1815 | ASTE1 | 2 |
| 300 | PSMB8-AS1 | 1 | 1056 | ATP6AP2 | 1 | 1816 | RLF | 2 |
| 301 | ABRACL | 1 | 1057 | LINC00847 | 1 | 1817 | SYTL3 | 2 |
| 302 | EPS8 | 1 | 1058 | PIM1 | 1 | 1818 | AL731571.1 | 2 |
| 303 | MPHOSPH8 | 1 | 1059 | SLAMF6 | 1 | 1819 | BAIAP2 | 2 |
| 304 | NCR1 | 1 | 1060 | PPM1G | 1 | 1820 | CKAP2 | 2 |
| 305 | RCSD1 | 1 | 1061 | UBASH3B | 1 | 1821 | HNRNPU | 2 |
| 306 | GNPTAB | 1 | 1062 | TOMM40 | 1 | 1822 | TUFT1 | 2 |
| 307 | DRAP1 | 1 | 1063 | LINC00476 | 1 | 1823 | PPP1R13B | 2 |
| 308 | TMBIM4 | 1 | 1064 | CRELD2 | 1 | 1824 | EIF4A1 | 2 |
| 309 | STMP1 | 1 | 1065 | GART | 1 | 1825 | ZRANB1 | 2 |
| 310 | COX7A2L | 1 | 1066 | TMEM161B | 1 | 1826 | SREBF1 | 2 |
| 311 | H3F3A | 1 | 1067 | SRSF4 | 1 | 1827 | SLC45A4 | 2 |
| 312 | TRGC1 | 1 | 1068 | ERV3-1 | 1 | 1828 | HP | 2 |
| 313 | KLRC4 | 1 | 1069 | MX1 | 1 | 1829 | RBMX | 2 |
| 314 | LAMTOR2 | 1 | 1070 | Z93930.2 | 1 | 1830 | HIST1H3B | 2 |
| 315 | RPP38 | 1 | 1071 | PSMD3 | 1 | 1831 | MED31 | 2 |
| 316 | MRPL51 | 1 | 1072 | QTRT1 | 1 | 1832 | STRAP | 2 |
| 317 | RAD23A | 1 | 1073 | ZNF429 | 1 | 1833 | UBE2H | 2 |
| 318 | WAS | 1 | 1074 | C11orf71 | 1 | 1834 | RPP21 | 2 |
| 319 | TXN2 | 1 | 1075 | MFNG | 1 | 1835 | WAC | 2 |
| 320 | PNN | 1 | 1076 | ARHGEF1 | 1 | 1836 | APOA2 | 2 |
| 321 | PARP1 | 1 | 1077 | SEPHS2 | 1 | 1837 | PHF1 | 2 |
| 322 | TMEM14B | 1 | 1078 | SLC16A1-A | 1 | 1838 | SHOC2 | 2 |
| 323 | COX5B | 1 | 1079 | DEF6 | 1 | 1839 | ZNF165 | 2 |
| 324 | GFI1 | 1 | 1080 | TTC13 | 1 | 1840 | CXCL8 | 2 |
| 325 | ZNF844 | 1 | 1081 | NDUFB5 | 1 | 1841 | YTHDC1 | 2 |
| 326 | SKAP1 | 1 | 1082 | PILRB | 1 | 1842 | HSPA5 | 2 |
| 327 | HMGN2 | 1 | 1083 | STX7 | 1 | 1843 | APOA1 | 2 |
| 328 | NDUFB3 | 1 | 1084 | LBH | 1 | 1844 | SMARCA5 | 2 |
| 329 | PPAN | 1 | 1085 | AAMP | 1 | 1845 | CDK5RAP1 | 2 |
| 330 | ARPC1B | 1 | 1086 | CASP2 | 1 | 1846 | CHKA | 2 |
| 331 | ITGB1BP1 | 1 | 1087 | ZNF83 | 1 | 1847 | ZPR1 | 2 |
| 332 | TUFM | 1 | 1088 | RPS19BP1 | 1 | 1848 | REV3L | 2 |
| 333 | PHB2 | 1 | 1089 | CIDEB | 1 | 1849 | MIR194-2H( | 2 |
| 334 | DYNLRB1 | 1 | 1090 | SCML4 | 1 | 1850 | AC104116.1 | 2 |
| 335 | ZNHIT6 | 1 | 1091 | RMRP | 1 | 1851 | EIF1AX | 2 |
| 336 | GOLGA8A | 1 | 1092 | COMMD10 | 1 | 1852 | GPM6B | 2 |
| 337 | NDUFAF3 | 1 | 1093 | POLDIP2 | 1 | 1853 | HIVEP2 | 2 |
| 338 | TBCB | 1 | 1094 | C6orf226 | 1 | 1854 | CCDC59 | 2 |

| | | | | | | | | |
|---|---|---|---|---|---|---|---|---|
| 339 | AC020911.2 | 1 | 1095 | UNG | 1 | 1855 | ZNF706 | 2 |
| 340 | CELF2 | 1 | 1096 | UQCRC2 | 1 | 1856 | ABCF1 | 2 |
| 341 | FADD | 1 | 1097 | COX18 | 1 | 1857 | DSCR9 | 2 |
| 342 | PYCARD-A | 1 | 1098 | GPANK1 | 1 | 1858 | SNRPB | 2 |
| 343 | AC005332.5 | 1 | 1099 | FOPNL | 1 | 1859 | ZNF10 | 2 |
| 344 | FAM45A | 1 | 1100 | DNAJC4 | 1 | 1860 | RAB1A | 2 |
| 345 | TIMM8B | 1 | 1101 | GPSM3 | 1 | 1861 | H2AFX | 2 |
| 346 | TRIM14 | 1 | 1102 | ANAPC4 | 1 | 1862 | AC026979.2 | 2 |
| 347 | ADM | 1 | 1103 | SNRNP200 | 1 | 1863 | ERO1B | 2 |
| 348 | THRAP3 | 1 | 1104 | AC027644.3 | 1 | 1864 | SOCS4 | 2 |
| 349 | P4HB | 1 | 1105 | LINC00891 | 1 | 1865 | KMT5C | 2 |
| 350 | SRP14 | 1 | 1106 | SPA17 | 1 | 1866 | ELOC | 2 |
| 351 | SNRNP70 | 1 | 1107 | AL136040.1 | 1 | 1867 | RAB5A | 2 |
| 352 | EDF1 | 1 | 1108 | HTATIP2 | 1 | 1868 | UQCRH | 2 |
| 353 | CAPN1 | 1 | 1109 | ARL17A | 1 | 1869 | EFNB2 | 2 |
| 354 | HPGD | 1 | 1110 | MCTS1 | 1 | 1870 | AC124798.1 | 2 |
| 355 | COPZ1 | 1 | 1111 | HDDC3 | 1 | 1871 | JMY | 2 |
| 356 | SMDT1 | 1 | 1112 | BRD9 | 1 | 1872 | FXR1 | 2 |
| 357 | ZAP70 | 1 | 1113 | UBE2G2 | 1 | 1873 | TCF7 | 2 |
| 358 | ERAP2 | 1 | 1114 | MYBL1 | 1 | 1874 | GPR35 | 2 |
| 359 | GIMAP6 | 1 | 1115 | RHOBTB3 | 1 | 1875 | WSB1 | 2 |
| 360 | LSM7 | 1 | 1116 | CTSA | 1 | 1876 | AHR | 2 |
| 361 | METTL9 | 1 | 1117 | NDUFA13 | 1 | 1877 | FEZ1 | 2 |
| 362 | NME3 | 1 | 1118 | HDAC10 | 1 | 1878 | CERK | 2 |
| 363 | ARPC5L | 1 | 1119 | LMF2 | 1 | 1879 | ATF3 | 3 |
| 364 | ZMAT2 | 1 | 1120 | FAAP20 | 1 | 1880 | CCL4 | 3 |
| 365 | TNFRSF1A | 1 | 1121 | SIRT7 | 1 | 1881 | CCL4L2 | 3 |
| 366 | NEMF | 1 | 1122 | CDK5RAP3 | 1 | 1882 | TNFAIP3 | 3 |
| 367 | ATP5F1B | 1 | 1123 | CASP6 | 1 | 1883 | GRASP | 3 |
| 368 | POLR2E | 1 | 1124 | MRPL52 | 1 | 1884 | KLF2 | 3 |
| 369 | SSR4 | 1 | 1125 | GPR174 | 1 | 1885 | CREM | 3 |
| 370 | TMBIM6 | 1 | 1126 | ZCRB1 | 1 | 1886 | NFE2L2 | 3 |
| 371 | FNBP4 | 1 | 1127 | USP48 | 1 | 1887 | GNLY | 3 |
| 372 | GUK1 | 1 | 1128 | TRIM13 | 1 | 1888 | KLF6 | 3 |
| 373 | CYB5D2 | 1 | 1129 | AZI2 | 1 | 1889 | PTGER4 | 3 |
| 374 | COA3 | 1 | 1130 | NDUFS3 | 1 | 1890 | TXNIP | 3 |
| 375 | MYDGF | 1 | 1131 | NIT2 | 1 | 1891 | NKG7 | 3 |
| 376 | TIA1 | 1 | 1132 | SRP9 | 1 | 1892 | BHLHE40 | 3 |
| 377 | COMMD6 | 1 | 1133 | CTSG | 1 | 1893 | PMAIP1 | 3 |
| 378 | PA2G4 | 1 | 1134 | YARS | 1 | 1894 | HLA-E | 3 |
| 379 | ZNF160 | 1 | 1135 | AC093462.1 | 1 | 1895 | CRIP1 | 3 |
| 380 | MDFIC | 1 | 1136 | PDCD2 | 1 | 1896 | ISG20 | 3 |
| 381 | SNX10 | 1 | 1137 | RIPK3 | 1 | 1897 | RPS4Y1 | 3 |
| 382 | SSH2 | 1 | 1138 | AKAP12 | 1 | 1898 | CD3E | 3 |
| 383 | PPP1R7 | 1 | 1139 | OGA | 1 | 1899 | ZFP36L2 | 3 |
| 384 | TALDO1 | 1 | 1140 | SNRPC | 1 | 1900 | ANXA1 | 3 |
| 385 | CTDSP1 | 1 | 1141 | PDLIM2 | 1 | 1901 | PER1 | 3 |
| 386 | 7-Sep | 1 | 1142 | PTPN18 | 1 | 1902 | SQSTM1 | 3 |
| 387 | ZBTB38 | 1 | 1143 | TRNAU1AP | 1 | 1903 | RPS3A | 3 |
| 388 | TRNT1 | 1 | 1144 | AC245014.3 | 1 | 1904 | PRF1 | 3 |
| 389 | ATP5MG | 1 | 1145 | ORMDL1 | 1 | 1905 | FTH1 | 3 |
| 390 | IQGAP1 | 1 | 1146 | TRIP11 | 1 | 1906 | RGS2 | 3 |
| 391 | KLHL6 | 1 | 1147 | EIF3I | 1 | 1907 | CYBA | 3 |
| 392 | PHB | 1 | 1148 | SLA2 | 1 | 1908 | IL32 | 3 |
| 393 | BLOC1S1 | 1 | 1149 | AC142472.1 | 1 | 1909 | RPL14 | 3 |
| 394 | DMAC1 | 1 | 1150 | ADAM28 | 1 | 1910 | MT-CYB | 3 |
| 395 | ORAI2 | 1 | 1151 | ARFGAP2 | 1 | 1911 | HLA-C | 3 |
| 396 | ATF6B | 1 | 1152 | C9orf64 | 1 | 1912 | RILPL2 | 3 |
| 397 | ICAM3 | 1 | 1153 | DDX43 | 1 | 1913 | CYTIP | 3 |
| 398 | CXorf38 | 1 | 1154 | NUDT16L1 | 1 | 1914 | VIM | 3 |
| 399 | EIF3J-DT | 1 | 1155 | OTUB1 | 1 | 1915 | SH3BGRL3 | 3 |
| 400 | NAA10 | 1 | 1156 | PGP | 1 | 1916 | TAGLN2 | 3 |
| 401 | SF3B2 | 1 | 1157 | PIGS | 1 | 1917 | BRD2 | 3 |
| 402 | IFI27L2 | 1 | 1158 | PPOX | 1 | 1918 | GPR65 | 3 |
| 403 | TADA3 | 1 | 1159 | ST3GAL1 | 1 | 1919 | IER3 | 3 |
| 404 | NUCB1 | 1 | 1160 | C8orf33 | 1 | 1920 | CTSW | 3 |
| 405 | ATRAID | 1 | 1161 | ENOPH1 | 1 | 1921 | PIM3 | 3 |
| 406 | BANF1 | 1 | 1162 | GTF2A2 | 1 | 1922 | GZMH | 3 |

| | | | | | | | | |
|---|---|---|---|---|---|---|---|---|
| 407 | GPAA1 | 1 | 1163 | SNX14 | 1 | 1923 | HSPB1 | 3 |
| 408 | UQCRQ | 1 | 1164 | ARFIP2 | 1 | 1924 | RPS4X | 3 |
| 409 | DGCR6L | 1 | 1165 | CTBS | 1 | 1925 | RUNX3 | 3 |
| 410 | NDUFB8 | 1 | 1166 | RAD9A | 1 | 1926 | SAT1 | 3 |
| 411 | CMTM3 | 1 | 1167 | POP4 | 1 | 1927 | LMNA | 3 |
| 412 | FAM50A | 1 | 1168 | MUS81 | 1 | 1928 | CD2 | 3 |
| 413 | RANGRF | 1 | 1169 | CHMP1A | 1 | 1929 | CDKN1A | 3 |
| 414 | HP1BP3 | 1 | 1170 | URI1 | 1 | 1930 | IVNS1ABP | 3 |
| 415 | CISD3 | 1 | 1171 | ZNF253 | 1 | 1931 | NEAT1 | 3 |
| 416 | TMEM223 | 1 | 1172 | RTL8A | 1 | 1932 | CD55 | 3 |
| 417 | HNRNPA3 | 1 | 1173 | LSM8 | 1 | 1933 | PHLDA1 | 3 |
| 418 | ILK | 1 | 1174 | DPYSL2 | 1 | 1934 | PRDM1 | 3 |
| 419 | ANKRD36 | 1 | 1175 | CSRP1 | 1 | 1935 | TMSB10 | 3 |
| 420 | PCSK7 | 1 | 1176 | ZDHHC12 | 1 | 1936 | MTRNR2L1 | 3 |
| 421 | POLR3GL | 1 | 1177 | AP001462.1 | 1 | 1937 | UAP1 | 3 |
| 422 | ITM2B | 1 | 1178 | ATXN7L3B | 1 | 1938 | ITGB7 | 3 |
| 423 | NDUFA12 | 1 | 1179 | PPP1R3D | 1 | 1939 | RPL7A | 3 |
| 424 | CHURC1 | 1 | 1180 | CLASP1 | 1 | 1940 | MT-CO2 | 3 |
| 425 | CYBC1 | 1 | 1181 | ATL3 | 1 | 1941 | UCP2 | 3 |
| 426 | PTPN12 | 1 | 1182 | KDELR1 | 1 | 1942 | RPS21 | 3 |
| 427 | CCNL2 | 1 | 1183 | NSUN5 | 1 | 1943 | CD3D | 3 |
| 428 | XRCC6 | 1 | 1184 | R3HCC1 | 1 | 1944 | PPP1R10 | 3 |
| 429 | YPEL1 | 1 | 1185 | NUDCD3 | 1 | 1945 | LITAF | 3 |
| 430 | RAB30-AS1 | 1 | 1186 | LIG1 | 1 | 1946 | RPL28 | 3 |
| 431 | SCP2 | 1 | 1187 | CLINT1 | 1 | 1947 | CARD16 | 3 |
| 432 | NDUFS2 | 1 | 1188 | SDHD | 1 | 1948 | TMEM173 | 3 |
| 433 | RAP1GDS1 | 1 | 1189 | PCNX1 | 1 | 1949 | SLFN11 | 3 |
| 434 | AC083973.1 | 1 | 1190 | HGH1 | 1 | 1950 | RPS5 | 3 |
| 435 | VPS28 | 1 | 1191 | ERGIC2 | 1 | 1951 | NEU1 | 3 |
| 436 | B4GALT4 | 1 | 1192 | SF3A3 | 1 | 1952 | RNF19A | 3 |
| 437 | MRPL57 | 1 | 1193 | UGP2 | 1 | 1953 | RPL21 | 3 |
| 438 | ZRANB2 | 1 | 1194 | KIN | 1 | 1954 | TNF | 3 |
| 439 | UROS | 1 | 1195 | YIF1A | 1 | 1955 | SH3BP5 | 3 |
| 440 | ARHGEF9 | 1 | 1196 | RNPEP | 1 | 1956 | HOPX | 3 |
| 441 | GATAD1 | 1 | 1197 | UBE2L3 | 1 | 1957 | KLF13 | 3 |
| 442 | PSMA1 | 1 | 1198 | RPUSD3 | 1 | 1958 | GNG2 | 3 |
| 443 | TXN | 1 | 1199 | TMEM183A | 1 | 1959 | S100A4 | 3 |
| 444 | ATP5PD | 1 | 1200 | ASCC1 | 1 | 1960 | CST7 | 3 |
| 445 | ELANE | 1 | 1201 | OARD1 | 1 | 1961 | CD3G | 3 |
| 446 | ZNHIT1 | 1 | 1202 | COA6 | 1 | 1962 | RPS27 | 3 |
| 447 | FKBP1A | 1 | 1203 | TBC1D1 | 1 | 1963 | B2M | 3 |
| 448 | TTC14 | 1 | 1204 | VMP1 | 1 | 1964 | SPON2 | 3 |
| 449 | PSMD8 | 1 | 1205 | DNPEP | 1 | 1965 | AES | 3 |
| 450 | LSM10 | 1 | 1206 | PRPSAP2 | 1 | 1966 | TLE4 | 3 |
| 451 | NDUFV1 | 1 | 1207 | YIPF2 | 1 | 1967 | RPS14 | 3 |
| 452 | RBCK1 | 1 | 1208 | ZNF701 | 1 | 1968 | RPS9 | 3 |
| 453 | GCHFR | 1 | 1209 | ARHGAP30 | 1 | 1969 | TNFRSF18 | 3 |
| 454 | BCL7C | 1 | 1210 | ZNF708 | 1 | 1970 | RPL39 | 3 |
| 455 | B3GALT4 | 1 | 1211 | DBI | 1 | 1971 | ANXA2 | 3 |
| 456 | COX4I1 | 1 | 1212 | NHLRC2 | 1 | 1972 | COQ10B | 3 |
| 457 | TMEM101 | 1 | 1213 | LEPROT | 1 | 1973 | MT-CO1 | 3 |
| 458 | CREBZF | 1 | 1214 | LINC00685 | 1 | 1974 | ADGRG1 | 3 |
| 459 | CCM2 | 1 | 1215 | LRPAP1 | 1 | 1975 | RPSA | 3 |
| 460 | SNF8 | 1 | 1216 | C18orf21 | 1 | 1976 | MLF1 | 3 |
| 461 | NDUFS6 | 1 | 1217 | EIF2B1 | 1 | 1977 | COX7C | 3 |
| 462 | GYG1 | 1 | 1218 | SMARCE1 | 1 | 1978 | SNRPD2 | 3 |
| 463 | GPS1 | 1 | 1219 | AL137077.2 | 1 | 1979 | TNFRSF1B | 3 |
| 464 | NHP2 | 1 | 1220 | PRPF38B | 1 | 1980 | BAZ1A | 3 |
| 465 | HNRNPA2E | 1 | 1221 | CEBPZOS | 1 | 1981 | NDUFB7 | 3 |
| 466 | MRPL41 | 1 | 1222 | MPPE1 | 1 | 1982 | S100A6 | 3 |
| 467 | TAPBP | 1 | 1223 | MRPL49 | 1 | 1983 | RPS3 | 3 |
| 468 | CPNE1 | 1 | 1224 | IWS1 | 1 | 1984 | AC091271.1 | 3 |
| 469 | SLC25A11 | 1 | 1225 | YPEL3 | 1 | 1985 | EMP3 | 3 |
| 470 | AP2S1 | 1 | 1226 | ARIH2OS | 1 | 1986 | BX284668.6 | 3 |
| 471 | MAGED2 | 1 | 1227 | RBM6 | 1 | 1987 | FCGR3A | 3 |
| 472 | TMEM59 | 1 | 1228 | PSMD13 | 1 | 1988 | MRPS6 | 3 |
| 473 | DPH7 | 1 | 1229 | RTCB | 1 | 1989 | SLC5A3 | 3 |
| 474 | ANAPC16 | 1 | 1230 | CRELD1 | 1 | 1990 | MBP | 3 |

| | | | | | | | | |
|---|---|---|---|---|---|---|---|---|
| 475 | ISCU | 1 | 1231 | CPT2 | 1 | 1991 | PTPN6 | 3 |
| 476 | P2RX5 | 1 | 1232 | PSME3 | 1 | 1992 | AC087239.1 | 3 |
| 477 | EAPP | 1 | 1233 | SNX3 | 1 | 1993 | MIR22HG | 3 |
| 478 | HIST1H2BH | 1 | 1234 | ZNF688 | 1 | 1994 | RPS15 | 3 |
| 479 | NDUFA11 | 1 | 1235 | TMEM9B | 1 | 1995 | RPL13 | 3 |
| 480 | PGLS | 1 | 1236 | OCIAD2 | 1 | 1996 | BAG3 | 3 |
| 481 | NCBP2-AS2 | 1 | 1237 | EIF2B2 | 1 | 1997 | ID3 | 3 |
| 482 | ARHGAP4 | 1 | 1238 | KLHL23 | 1 | 1998 | RPS25 | 3 |
| 483 | MRPS18B | 1 | 1239 | RNF181 | 1 | 1999 | STK4 | 3 |
| 484 | COPS6 | 1 | 1240 | NDUFB11 | 1 | 2000 | RPS29 | 3 |
| 485 | MZB1 | 1 | 1241 | IFRD2 | 1 | 2001 | RPL18A | 3 |
| 486 | NCK2 | 1 | 1242 | PAFAH2 | 1 | 2002 | RPL19 | 3 |
| 487 | RPN2 | 1 | 1243 | GTF3C5 | 1 | 2003 | LDHB | 3 |
| 488 | UBL4A | 1 | 1244 | NFATC2IP | 1 | 2004 | SMAP2 | 3 |
| 489 | MAPK13 | 1 | 1245 | ASB8 | 1 | 2005 | BIN1 | 3 |
| 490 | S100PBP | 1 | 1246 | GEMIN6 | 1 | 2006 | LIMD2 | 3 |
| 491 | TSPAN31 | 1 | 1247 | RGPD2 | 1 | 2007 | IL7R | 3 |
| 492 | TMEM219 | 1 | 1248 | SCAPER | 1 | 2008 | OSM | 3 |
| 493 | SERPINB6 | 1 | 1249 | MTERF3 | 1 | 2009 | AVPI1 | 3 |
| 494 | SDHC | 1 | 1250 | TTLL3 | 1 | 2010 | RPL37 | 3 |
| 495 | FYB1 | 1 | 1251 | C16orf58 | 1 | 2011 | RPL29 | 3 |
| 496 | NDUFS8 | 1 | 1252 | WDR61 | 1 | 2012 | CD8A | 3 |
| 497 | CCNDBP1 | 1 | 1253 | COMMD8 | 1 | 2013 | CD8B | 3 |
| 498 | FBXO2 | 1 | 1254 | AL606760.3 | 1 | 2014 | RPS15A | 3 |
| 499 | PPIH | 1 | 1255 | URM1 | 1 | 2015 | RPL6 | 3 |
| 500 | MCRIP1 | 1 | 1256 | AC064807.1 | 1 | 2016 | TNFRSF9 | 3 |
| 501 | KLRB1 | 1 | 1257 | DDX28 | 1 | 2017 | CDC42SE1 | 3 |
| 502 | MRPS23 | 1 | 1258 | METTL18 | 1 | 2018 | DUSP8 | 3 |
| 503 | PDIA3 | 1 | 1259 | RNF130 | 1 | 2019 | GZMB | 3 |
| 504 | RPE | 1 | 1260 | PFDN2 | 1 | 2020 | LDLR | 3 |
| 505 | C11orf68 | 1 | 1261 | RAC1 | 1 | 2021 | RPL23A | 3 |
| 506 | TRAPPC6A | 1 | 1262 | NECAP2 | 1 | 2022 | ARL4C | 3 |
| 507 | COX7A2 | 1 | 1263 | MBIP | 1 | 2023 | ERRFI1 | 3 |
| 508 | CTSZ | 1 | 1264 | SRSF1 | 1 | 2024 | ITPRIP | 3 |
| 509 | ZNF224 | 1 | 1265 | PPM1M | 1 | 2025 | CCND3 | 3 |
| 510 | AKR1B1 | 1 | 1266 | SCCPDH | 1 | 2026 | MT-ND1 | 3 |
| 511 | SRI | 1 | 1267 | FUCA1 | 1 | 2027 | RACK1 | 3 |
| 512 | AGTRAP | 1 | 1268 | VPS4B | 1 | 2028 | HLA-DRB1 | 3 |
| 513 | HNRNPR | 1 | 1269 | NKILA | 1 | 2029 | TTC38 | 3 |
| 514 | ARFRP1 | 1 | 1270 | RAB37 | 1 | 2030 | SOX4 | 3 |
| 515 | MRPL23 | 1 | 1271 | CENPX | 1 | 2031 | RPL10 | 3 |
| 516 | MED7 | 1 | 1272 | ESF1 | 1 | 2032 | RABAC1 | 3 |
| 517 | LAMP1 | 1 | 1273 | MRPS14 | 1 | 2033 | RPS28 | 3 |
| 518 | NDUFA9 | 1 | 1274 | COA5 | 1 | 2034 | TPST2 | 3 |
| 519 | NCAM1 | 1 | 1275 | ANXA11 | 1 | 2035 | HSH2D | 3 |
| 520 | C9orf78 | 1 | 1276 | ACP1 | 1 | 2036 | RPL35A | 3 |
| 521 | GFOD1 | 1 | 1277 | CTSC | 1 | 2037 | RPL27A | 3 |
| 522 | SRSF11 | 1 | 1278 | RWDD1 | 1 | 2038 | HEXIM1 | 3 |
| 523 | CHMP4A | 1 | 1279 | MZF1-AS1 | 1 | 2039 | TMEM71 | 3 |
| 524 | LENG8 | 1 | 1280 | AL031708.1 | 1 | 2040 | CASP8 | 3 |
| 525 | EPSTI1 | 1 | 1281 | MED8 | 1 | 2041 | OAT | 3 |
| 526 | SMIM7 | 1 | 1282 | SNRPA1 | 1 | 2042 | RPL30 | 3 |
| 527 | CNBP | 1 | 1283 | MIR4435-2H | 1 | 2043 | KLRG1 | 3 |
| 528 | SELENOH | 1 | 1284 | TNFSF10 | 1 | 2044 | PPDPF | 3 |
| 529 | PRSS21 | 1 | 1285 | ELP5 | 1 | 2045 | JAK1 | 3 |
| 530 | METTL17 | 1 | 1286 | PPP1R12A | 1 | 2046 | ATP2B1-AS | 3 |
| 531 | LINC02256 | 1 | 1287 | AC137767.1 | 1 | 2047 | RPS24 | 3 |
| 532 | ACAA2 | 1 | 1288 | ABHD11 | 1 | 2048 | RPL34 | 3 |
| 533 | GRK2 | 1 | 1289 | C22orf39 | 1 | 2049 | ZC3HAV1 | 3 |
| 534 | IGHG4 | 1 | 1290 | PIN1 | 1 | 2050 | DDIT3 | 3 |
| 535 | PLA2G16 | 1 | 1291 | AP4B1 | 1 | 2051 | MT-ATP6 | 3 |
| 536 | DPM2 | 1 | 1292 | SELENOT | 1 | 2052 | SELPLG | 3 |
| 537 | PSMC5 | 1 | 1293 | ELF2 | 1 | 2053 | ARPC2 | 3 |
| 538 | TNFSF12 | 1 | 1294 | SPNS3 | 1 | 2054 | CD52 | 3 |
| 539 | REEP5 | 1 | 1295 | LINC02001 | 1 | 2055 | SAMHD1 | 3 |
| 540 | DCAF7 | 1 | 1296 | C2orf68 | 1 | 2056 | RASGRP2 | 3 |
| 541 | OSTC | 1 | 1297 | KEAP1 | 1 | 2057 | MALAT1 | 3 |
| 542 | RCCD1 | 1 | 1298 | CHKB | 1 | 2058 | CAPG | 3 |

| | | | | | | | | |
|---|---|---|---|---|---|---|---|---|
| 543 | PYURF | 1 | 1299 | DNASE1 | 1 | 2059 | PRDX2 | 3 |
| 544 | FAM173A | 1 | 1300 | SSU72 | 1 | 2060 | NCR3 | 3 |
| 545 | IAH1 | 1 | 1301 | B3GAT3 | 1 | 2061 | TTTY15 | 3 |
| 546 | AC025164.1 | 1 | 1302 | NME6 | 1 | 2062 | RPLP2 | 3 |
| 547 | GON4L | 1 | 1303 | AREG | 2 | 2063 | LGALS1 | 3 |
| 548 | AL355472.1 | 1 | 1304 | ATP1B3 | 2 | 2064 | HIF1A | 3 |
| 549 | APOBEC3C | 1 | 1305 | CD69 | 2 | 2065 | PPIA | 3 |
| 550 | RPL7L1 | 1 | 1306 | DNAJA1 | 2 | 2066 | XBP1 | 3 |
| 551 | SHISA5 | 1 | 1307 | DNAJB1 | 2 | 2067 | SGK1 | 3 |
| 552 | CD151 | 1 | 1308 | DNAJB6 | 2 | 2068 | RPS16 | 3 |
| 553 | ACP5 | 1 | 1309 | DUSP1 | 2 | 2069 | MT-ND4 | 3 |
| 554 | TLE1 | 1 | 1310 | DUSP2 | 2 | 2070 | RPS19 | 3 |
| 555 | AC245297.3 | 1 | 1311 | FAM177A1 | 2 | 2071 | UPP1 | 3 |
| 556 | SUPT16H | 1 | 1312 | FOSB | 2 | 2072 | RPL10A | 3 |
| 557 | HENMT1 | 1 | 1313 | HSP90AA1 | 2 | 2073 | RPS18 | 3 |
| 558 | CDK2AP2 | 1 | 1314 | HSP90AB1 | 2 | 2074 | SYNE2 | 3 |
| 559 | IL16 | 1 | 1315 | HSPA1A | 2 | 2075 | LINC00861 | 3 |
| 560 | TSR2 | 1 | 1316 | HSPA1B | 2 | 2076 | ULBP2 | 3 |
| 561 | PRDX6 | 1 | 1317 | HSPA8 | 2 | 2077 | SOD1 | 3 |
| 562 | TRIM73 | 1 | 1318 | HSPD1 | 2 | 2078 | GLA | 3 |
| 563 | COX17 | 1 | 1319 | HSPE1 | 2 | 2079 | GBP5 | 3 |
| 564 | RCN2 | 1 | 1320 | HSPH1 | 2 | 2080 | TRBC1 | 3 |
| 565 | EIF2S3 | 1 | 1321 | IRF1 | 2 | 2081 | Z93241.1 | 3 |
| 566 | CRACR2B | 1 | 1322 | JUN | 2 | 2082 | IQCN | 3 |
| 567 | PSMD4 | 1 | 1323 | JUNB | 2 | 2083 | NACA | 3 |
| 568 | FGR | 1 | 1324 | JUND | 2 | 2084 | CLEC2B | 3 |
| 569 | TMEM140 | 1 | 1325 | METRNL | 2 | 2085 | C12orf57 | 3 |
| 570 | P2RY11 | 1 | 1326 | NFKBIA | 2 | 2086 | MT-CO3 | 3 |
| 571 | NUTM2B-A | 1 | 1327 | NR4A1 | 2 | 2087 | SUN2 | 3 |
| 572 | PGAM1 | 1 | 1328 | NR4A2 | 2 | 2088 | MT2A | 3 |
| 573 | SURF6 | 1 | 1329 | PABPC1 | 2 | 2089 | SSR2 | 3 |
| 574 | NUBP2 | 1 | 1330 | PDE4B | 2 | 2090 | UTY | 3 |
| 575 | DGUOK | 1 | 1331 | PPP1R15A | 2 | 2091 | CLCF1 | 3 |
| 576 | TAF8 | 1 | 1332 | REL | 2 | 2092 | BIN2 | 3 |
| 577 | IFI44 | 1 | 1333 | RGCC | 2 | 2093 | GLIPR2 | 3 |
| 578 | COA4 | 1 | 1334 | TIPARP | 2 | 2094 | RPS8 | 3 |
| 579 | HMGB2 | 1 | 1335 | TWISTNB | 2 | 2095 | LCK | 3 |
| 580 | RFC1 | 1 | 1336 | UBB | 2 | 2096 | S1PR5 | 3 |
| 581 | KRT10 | 1 | 1337 | YPEL5 | 2 | 2097 | RPL11 | 3 |
| 582 | KRT81 | 1 | 1338 | ZFP36 | 2 | 2098 | RPS23 | 3 |
| 583 | SUGP2 | 1 | 1339 | ZNF331 | 2 | 2099 | RPL37A | 3 |
| 584 | IARS2 | 1 | 1340 | CD83 | 2 | 2100 | UBA52 | 3 |
| 585 | LAT | 1 | 1341 | CHMP1B | 2 | 2101 | MTHFD2 | 3 |
| 586 | TRABD | 1 | 1342 | HSPA6 | 2 | 2102 | MT1X | 3 |
| 587 | AC005837.1 | 1 | 1343 | UBC | 2 | 2103 | AL450998.2 | 3 |
| 588 | SSNA1 | 1 | 1344 | CEMIP2 | 2 | 2104 | KLF9 | 3 |
| 589 | PSMA7 | 1 | 1345 | NFKBIZ | 2 | 2105 | RPL36AL | 3 |
| 590 | C19orf25 | 1 | 1346 | ARL5B | 2 | 2106 | KMT2E-AS1 | 3 |
| 591 | THUMPD2 | 1 | 1347 | ARL4A | 2 | 2107 | CDC42EP3 | 3 |
| 592 | FIBP | 1 | 1348 | KDM6B | 2 | 2108 | MAPRE2 | 3 |
| 593 | GSDMD | 1 | 1349 | CCL3 | 2 | 2109 | TRGV4 | 3 |
| 594 | SNRPA | 1 | 1350 | BTG2 | 2 | 2110 | RPL26 | 3 |
| 595 | TPM3 | 1 | 1351 | MCL1 | 2 | 2111 | BPGM | 3 |
| 596 | COX6A1 | 1 | 1352 | NR4A3 | 2 | 2112 | CD300A | 3 |
| 597 | SPAG7 | 1 | 1353 | CSRNP1 | 2 | 2113 | AC074044.1 | 3 |
| 598 | CYC1 | 1 | 1354 | ICAM1 | 2 | 2114 | RPL41 | 3 |
| 599 | PWWP2A | 1 | 1355 | SELENOK | 2 | 2115 | SYTL1 | 3 |
| 600 | HEBP2 | 1 | 1356 | TUBA4A | 2 | 2116 | ADRB2 | 3 |
| 601 | MDM4 | 1 | 1357 | TNFSF9 | 2 | 2117 | MPV17 | 3 |
| 602 | NKTR | 1 | 1358 | CXCR4 | 2 | 2118 | RPS27A | 3 |
| 603 | PNKP | 1 | 1359 | UBE2S | 2 | 2119 | HNRNPL | 3 |
| 604 | GUSB | 1 | 1360 | CHORDC1 | 2 | 2120 | AL139274.2 | 3 |
| 605 | TSPO | 1 | 1361 | TSC22D3 | 2 | 2121 | INTS6-AS1 | 3 |
| 606 | CCS | 1 | 1362 | IFNG | 2 | 2122 | PRMT5-AS1 | 3 |
| 607 | MRPL43 | 1 | 1363 | MAFF | 2 | 2123 | RPL38 | 3 |
| 608 | NDUFAF8 | 1 | 1364 | JMJD6 | 2 | 2124 | HLA-DRA | 3 |
| 609 | DUSP23 | 1 | 1365 | H2AFZ | 2 | 2125 | KLRC2 | 3 |
| 610 | ALDH9A1 | 1 | 1366 | IFRD1 | 2 | 2126 | MT-ND2 | 3 |

| | | | | | | | | |
|---|---|---|---|---|---|---|---|---|
| 611 | PDIA6 | 1 | 1367 | DDX3Y | 2 | 2127 | MFSD10 | 3 |
| 612 | ZNF91 | 1 | 1368 | FOS | 2 | 2128 | MT-ND5 | 3 |
| 613 | CSNK2B | 1 | 1369 | NFKB1 | 2 | 2129 | FTL | 3 |
| 614 | ZNF652 | 1 | 1370 | ZFAND5 | 2 | 2130 | RPL9 | 3 |
| 615 | BCO2 | 1 | 1371 | SPTY2D1 | 2 | 2131 | SNHG25 | 3 |
| 616 | CAPNS1 | 1 | 1372 | RGS1 | 2 | 2132 | TOMM7 | 3 |
| 617 | SLC25A45 | 1 | 1373 | IDI1 | 2 | 2133 | YWHAQ | 3 |
| 618 | DHRS3 | 1 | 1374 | SERTAD1 | 2 | 2134 | HLA-DPB1 | 3 |
| 619 | ANKRD36C | 1 | 1375 | FOSL2 | 2 | 2135 | ADAP1 | 3 |
| 620 | CORO1B | 1 | 1376 | SKIL | 2 | 2136 | TRAC | 3 |
| 621 | PSMD7 | 1 | 1377 | CCNH | 2 | 2137 | MT-ATP8 | 3 |
| 622 | TP53TG1 | 1 | 1378 | RASGEF1B | 2 | 2138 | SLA | 3 |
| 623 | TSTA3 | 1 | 1379 | ELF1 | 2 | 2139 | LACTB | 3 |
| 624 | UQCC3 | 1 | 1380 | ZC3H12A | 2 | 2140 | PTPN4 | 3 |
| 625 | TMEM273 | 1 | 1381 | CACYBP | 2 | 2141 | SP140 | 3 |
| 626 | TMED2 | 1 | 1382 | VPS37B | 2 | 2142 | B4GALT1 | 3 |
| 627 | HDAC3 | 1 | 1383 | CKS2 | 2 | 2143 | ABHD17A | 3 |
| 628 | USP28 | 1 | 1384 | PPP1R2 | 2 | 2144 | PFDN5 | 3 |
| 629 | IDH2 | 1 | 1385 | MYADM | 2 | 2145 | RPL36A | 3 |
| 630 | SELENOF | 1 | 1386 | MRPL18 | 2 | 2146 | LYST | 3 |
| 631 | PYCR2 | 1 | 1387 | DUSP5 | 2 | 2147 | STAT3 | 3 |
| 632 | DPP7 | 1 | 1388 | RANBP2 | 2 | 2148 | AHNAK | 3 |
| 633 | CDK5R1 | 1 | 1389 | PIK3R1 | 2 | 2149 | RGS19 | 3 |
| 634 | 2-Mar | 1 | 1390 | ARF4 | 2 | 2150 | AC023509.4 | 3 |
| 635 | REX1BD | 1 | 1391 | ELL2 | 2 | 2151 | NFKBIB | 3 |
| 636 | DDX17 | 1 | 1392 | EGR1 | 2 | 2152 | MLLT11 | 3 |
| 637 | SH2D1A | 1 | 1393 | DYNLL1 | 2 | 2153 | U2AF1 | 3 |
| 638 | TRIM44 | 1 | 1394 | EIF1 | 2 | 2154 | AKAP13 | 3 |
| 639 | SNRPE | 1 | 1395 | H3F3B | 2 | 2155 | GLRX | 3 |
| 640 | MDH1 | 1 | 1396 | XCL2 | 2 | 2156 | MTRNR2L1 | 3 |
| 641 | PDHB | 1 | 1397 | IER5 | 2 | 2157 | DOK1 | 3 |
| 642 | GIMAP2 | 1 | 1398 | MAP3K8 | 2 | 2158 | ZFP36L1 | 3 |
| 643 | GLOD4 | 1 | 1399 | SCX | 2 | 2159 | RPS7 | 3 |
| 644 | LMAN2 | 1 | 1400 | SYAP1 | 2 | 2160 | AC009812.1 | 3 |
| 645 | CCR1 | 1 | 1401 | ID1 | 2 | 2161 | RPL36 | 3 |
| 646 | CNPY3 | 1 | 1402 | TPT1 | 2 | 2162 | AZU1 | 3 |
| 647 | OXNAD1 | 1 | 1403 | ANKRD37 | 2 | 2163 | CD37 | 3 |
| 648 | PFKL | 1 | 1404 | LDHA | 2 | 2164 | EIF3D | 3 |
| 649 | NDUFA8 | 1 | 1405 | NAF1 | 2 | 2165 | NDUFS7 | 3 |
| 650 | TSEN54 | 1 | 1406 | EIF4A3 | 2 | 2166 | CIB1 | 3 |
| 651 | WDR74 | 1 | 1407 | ZBTB1 | 2 | 2167 | GPR108 | 3 |
| 652 | HEXA | 1 | 1408 | GTF2B | 2 | 2168 | JAKMIP1 | 3 |
| 653 | POLM | 1 | 1409 | SRSF2 | 2 | 2169 | FGFBP2 | 3 |
| 654 | WASHC1 | 1 | 1410 | ETV3 | 2 | 2170 | TBRG1 | 3 |
| 655 | AKR7A2 | 1 | 1411 | NINJ1 | 2 | 2171 | MRPS21 | 3 |
| 656 | AIP | 1 | 1412 | SARAF | 2 | 2172 | RPL27 | 3 |
| 657 | ACADVL | 1 | 1413 | HES4 | 2 | 2173 | CAPN2 | 3 |
| 658 | JPX | 1 | 1414 | IRF8 | 2 | 2174 | ARHGEF37 | 3 |
| 659 | TMED10 | 1 | 1415 | PLIN2 | 2 | 2175 | RPS12 | 3 |
| 660 | TCEAL8 | 1 | 1416 | RAB8B | 2 | 2176 | ZMYND8 | 3 |
| 661 | CDK9 | 1 | 1417 | AC020916.1 | 2 | 2177 | ANP32B | 3 |
| 662 | CDK10 | 1 | 1418 | GNL3 | 2 | 2178 | ZBP1 | 3 |
| 663 | PRPF31 | 1 | 1419 | NXT1 | 2 | 2179 | PLK2 | 3 |
| 664 | LSM3 | 1 | 1420 | DNAJB4 | 2 | 2180 | SSBP1 | 3 |
| 665 | DUT | 1 | 1421 | RALGAPA1 | 2 | 2181 | SNAI1 | 3 |
| 666 | FAM118A | 1 | 1422 | PPP1R15B | 2 | 2182 | TXNL4A | 3 |
| 667 | CBLB | 1 | 1423 | SIK1 | 2 | 2183 | HAVCR2 | 3 |
| 668 | NDUFA1 | 1 | 1424 | FKBP4 | 2 | 2184 | HSPA2 | 3 |
| 669 | PDIA4 | 1 | 1425 | PRR7 | 2 | 2185 | CCDC173 | 3 |
| 670 | SCAMP1-A[illegible] | 1 | 1426 | ZFAND2A | 2 | 2186 | RALGDS | 3 |
| 671 | MRPL15 | 1 | 1427 | STX11 | 2 | 2187 | RPS11 | 3 |
| 672 | PIN4 | 1 | 1428 | PFKFB3 | 2 | 2188 | SCLT1 | 3 |
| 673 | JTB | 1 | 1429 | CCDC107 | 2 | 2189 | DAZAP2 | 3 |
| 674 | SMIM27 | 1 | 1430 | YWHAZ | 2 | 2190 | SAMD3 | 3 |
| 675 | SMIM20 | 1 | 1431 | PTMA | 2 | 2191 | ABHD5 | 3 |
| 676 | UBE2I | 1 | 1432 | CYCS | 2 | 2192 | S1PR4 | 3 |
| 677 | RNF135 | 1 | 1433 | BCL2A1 | 2 | 2193 | BCL11B | 3 |
| 678 | PEF1 | 1 | 1434 | IER2 | 2 | 2194 | MX2 | 3 |

| | | | | | | | | |
|---|---|---|---|---|---|---|---|---|
| 679 | CHMP6 | 1 | 1435 | GPBP1 | 2 | 2195 | RPS2 | 3 |
| 680 | WASHC3 | 1 | 1436 | EIF5 | 2 | 2196 | BTN3A2 | 3 |
| 681 | ANAPC11 | 1 | 1437 | ARIH1 | 2 | 2197 | DHRS7 | 3 |
| 682 | OGG1 | 1 | 1438 | ADGRE5 | 2 | 2198 | RND1 | 3 |
| 683 | LSM4 | 1 | 1439 | BCAS2 | 2 | 2199 | PYM1 | 3 |
| 684 | MYH9 | 1 | 1440 | EZR | 2 | 2200 | ANKRD9 | 3 |
| 685 | TCTA | 1 | 1441 | CALM1 | 2 | 2201 | HINT1 | 3 |
| 686 | N4BP2L2 | 1 | 1442 | AC044849.1 | 2 | 2202 | RYK | 3 |
| 687 | VASP | 1 | 1443 | NOCT | 2 | 2203 | POLR2A | 3 |
| 688 | PPP1R35 | 1 | 1444 | SDCBP | 2 | 2204 | AL356488.3 | 3 |
| 689 | YDJC | 1 | 1445 | NFIL3 | 2 | 2205 | CD99 | 3 |
| 690 | GADD45GI | 1 | 1446 | SRGN | 2 | 2206 | RPS6 | 3 |
| 691 | PET100 | 1 | 1447 | STK17B | 2 | 2207 | TSPAN32 | 3 |
| 692 | SLC2A4RG | 1 | 1448 | DCTN6 | 2 | 2208 | CRABP2 | 3 |
| 693 | FBXO22 | 1 | 1449 | TCP1 | 2 | 2209 | SLC38A2 | 3 |
| 694 | DCAF11 | 1 | 1450 | ETF1 | 2 | 2210 | AL451165.2 | 3 |
| 695 | PUF60 | 1 | 1451 | FAM53C | 2 | 2211 | ZNF276 | 3 |
| 696 | RINL | 1 | 1452 | DDX24 | 2 | 2212 | HNRNPA1 | 3 |
| 697 | CLSTN3 | 1 | 1453 | SLC7A5 | 2 | 2213 | PRDX1 | 3 |
| 698 | UHMK1 | 1 | 1454 | DDX5 | 2 | 2214 | POLG2 | 3 |
| 699 | ING4 | 1 | 1455 | NFKBID | 2 | 2215 | UBL3 | 3 |
| 700 | SELENOW | 1 | 1456 | AC016831.7 | 2 | 2216 | EHD4 | 3 |
| 701 | TYMP | 1 | 1457 | DDX3X | 2 | 2217 | RPL4 | 3 |
| 702 | NOP10 | 1 | 1458 | SOCS3 | 2 | 2218 | RIPOR2 | 3 |
| 703 | ATIC | 1 | 1459 | RBBP8 | 2 | 2219 | C12orf75 | 3 |
| 704 | SMC3 | 1 | 1460 | TSPYL2 | 2 | 2220 | MRPL16 | 3 |
| 705 | ZNF677 | 1 | 1461 | CHD1 | 2 | 2221 | PRRG2 | 3 |
| 706 | PSMB2 | 1 | 1462 | DEDD2 | 2 | 2222 | SELL | 3 |
| 707 | HLA-F | 1 | 1463 | BIRC3 | 2 | 2223 | TMEM238 | 3 |
| 708 | AC246785.3 | 1 | 1464 | HCG18 | 2 | 2224 | MXD1 | 3 |
| 709 | C22orf46 | 1 | 1465 | CCND2 | 2 | 2225 | FXYD5 | 3 |
| 710 | PDAP1 | 1 | 1466 | RELB | 2 | 2226 | SURF1 | 3 |
| 711 | AUP1 | 1 | 1467 | PTP4A1 | 2 | 2227 | FAM129A | 3 |
| 712 | DDT | 1 | 1468 | ODC1 | 2 | 2228 | RPL15 | 3 |
| 713 | NUDT18 | 1 | 1469 | BCL3 | 2 | 2229 | TAP1 | 3 |
| 714 | CCDC90B | 1 | 1470 | KMT2E | 2 | 2230 | TRAT1 | 3 |
| 715 | DCTN3 | 1 | 1471 | NEK1 | 2 | 2231 | XPA | 3 |
| 716 | RNF146 | 1 | 1472 | RRAD | 2 | 2232 | C7orf50 | 3 |
| 717 | NDUFB9 | 1 | 1473 | BZW1 | 2 | 2233 | HECA | 3 |
| 718 | ANKRD49 | 1 | 1474 | KLF10 | 2 | 2234 | AC106739.2 | 3 |
| 719 | COMMD7 | 1 | 1475 | MARCKSL1 | 2 | 2235 | AC104695.3 | 3 |
| 720 | NMT2 | 1 | 1476 | DNAJA4 | 2 | 2236 | RPL22 | 3 |
| 721 | NMUR1 | 1 | 1477 | ANKRD28 | 2 | 2237 | RPLP0 | 3 |
| 722 | AC008105.3 | 1 | 1478 | IFFO2 | 2 | 2238 | BAMBI | 3 |
| 723 | TRIM52 | 1 | 1479 | ERF | 2 | 2239 | LINC02446 | 3 |
| 724 | CCNT2 | 1 | 1480 | XCL1 | 2 | 2240 | C1QA | 3 |
| 725 | ELOVL6 | 1 | 1481 | BTG1 | 2 | 2241 | ZRSR2 | 3 |
| 726 | DDOST | 1 | 1482 | TAGAP | 2 | 2242 | TKT | 3 |
| 727 | ZNF43 | 1 | 1483 | CALM2 | 2 | 2243 | RPS13 | 3 |
| 728 | VAMP5 | 1 | 1484 | SBDS | 2 | 2244 | ZNF584 | 3 |
| 729 | AL135791.1 | 1 | 1485 | HIST1H4C | 2 | 2245 | FCGR3B | 3 |
| 730 | MRPL11 | 1 | 1486 | TENT5C | 2 | 2246 | TUBB2A | 3 |
| 731 | ARHGEF10 | 1 | 1487 | CSF2 | 2 | 2247 | HLA-DRB5 | 3 |
| 732 | CD96 | 1 | 1488 | CRTAM | 2 | 2248 | HAX1 | 3 |
| 733 | NMRAL1 | 1 | 1489 | SAMSN1 | 2 | 2249 | KLRC3 | 3 |
| 734 | GRAMD1C | 1 | 1490 | ZNF184 | 2 | 2250 | PCID2 | 3 |
| 735 | CINP | 1 | 1491 | YES1 | 2 | 2251 | RECQL | 3 |
| 736 | NDUFB4 | 1 | 1492 | NUFIP2 | 2 | 2252 | HMBOX1 | 3 |
| 737 | MRPL40 | 1 | 1493 | G3BP2 | 2 | 2253 | DIP2A | 3 |
| 738 | RNPEPL1 | 1 | 1494 | BRAF | 2 | 2254 | SLC27A3 | 3 |
| 739 | CCDC66 | 1 | 1495 | CCL3L1 | 2 | 2255 | GHDC | 3 |
| 740 | RGL4 | 1 | 1496 | NPM1 | 2 | 2256 | LINC02076 | 3 |
| 741 | ITGAE | 1 | 1497 | ZFY | 2 | 2257 | SNHG12 | 3 |
| 742 | NDUFC1 | 1 | 1498 | MIR155HG | 2 | 2258 | SLCO3A1 | 3 |
| 743 | XPO1 | 1 | 1499 | OTULIN | 2 | 2259 | CRBN | 3 |
| 744 | DCTN2 | 1 | 1500 | RBM8A | 2 | 2260 | NIPSNAP2 | 3 |
| 745 | UBLCP1 | 1 | 1501 | B3GNT7 | 2 | 2261 | IQGAP2 | 3 |
| 746 | TMX4 | 1 | 1502 | PRMT9 | 2 | 2262 | ARRB2 | 3 |

| | | |
|---|---|---|
| 747 | DCTD | 1 |
| 748 | SPG7 | 1 |
| 749 | FDPS | 1 |
| 750 | ANAPC15 | 1 |
| 751 | DENR | 1 |
| 752 | IFI6 | 1 |
| 753 | ZNF766 | 1 |
| 754 | AC112907.3 | 1 |
| 755 | MRPL17 | 1 |
| 756 | AUH | 1 |
| 1503 | CCNL1 | 2 |
| 1504 | TPM4 | 2 |
| 1505 | DOK2 | 2 |
| 1506 | RGS16 | 2 |
| 1507 | JOSD1 | 2 |
| 1508 | TUBB4B | 2 |
| 1509 | IDS | 2 |
| 1510 | RORA | 2 |
| 1511 | MAP1LC3B | 2 |
| 1512 | PNRC1 | 2 |
| 1513 | ERN1 | 2 |
| 1514 | RNF139 | 2 |
| 1515 | SMAD7 | 2 |
| 1516 | DENND4A | 2 |
| 2263 | RPL18 | 3 |
| 2264 | DTHD1 | 3 |
| 2265 | SP140L | 3 |
| 2266 | ERP29 | 3 |
| 2267 | CCDC25 | 3 |
| 2268 | ARHGEF2 | 3 |
| 2269 | AC018653.3 | 3 |
| 2270 | RUFY2 | 3 |
| 2271 | STOML2 | 3 |
| 2272 | CLK2 | 3 |
| 2273 | ILKAP | 3 |
| 2274 | SMCHD1 | 3 |

**Supplementary Table S3**

| Pro-inflammatory signature | Immune regulatory signature | Interferon responsed signature | Lipid metabolism signature |
|---|---|---|---|
| CCL3 | CD274 | ISG20 | ACP5 |
| CCL4 | CD40 | ISG15 | LPL |
| CCL20 | CD80 | IFI44L | TREM2 |
| CCL3L1 | CD86 | GBP1 | CCL18 |
| CCL4L2 | IDO1 | CASP1 | CTSB |
| CXCL1 | ICOSLG | CASP4 | CTSD |
| CXCL2 | IL10 | CXCL9 | CTSL |
| CXCL3 | TGFB1 | CXCL10 | FABP5 |
| CXCL5 | | CXCL11 | FABP4 |
| CXCL8 | | IFIT1 | ALOX5 AP |
| IL1B | | IFIT2 | |
| AREG | | IFIT3 | |
| EREG | | IFITM1 | |
| HBEGF | | IFITM3 | |

| Neutrophil maturation | Neutrophil chemotaxis | Phagocytosis | Type I interferon signaling pathway | Chemokine activity |
|---|---|---|---|---|
| SELPLG | BSG | A0A087WW49 | STAT2 | C5 |
| SAT1 | C1QBP | MYH9 | IFNAR1 | CCL1 |
| GRINA | C3AR1 | IGHV4-59 | CDC37 | CCL11 |
| CCL23 | C5AR1 | IGHV4-39 | H7C3V1 | CCL13 |
| CCL15 | C5AR2 | IGHV2-5 | PTPN6 | CCL14 |
| CCL14 | CAMK1D | IGHV3OR16-9 | IFNA8 | CCL15 |
| CEBPB | CCL1 | TREM2 | IFIT5 | CCL16 |
| ANXA2 | CCL11 | IGHV2-70 | TBK1 | CCL17 |
| GDA | CCL13 | IGKC | GBP2 | CCL18 |
| CLEC4D | CCL14 | TRBC1 | PTPN2 | CCL19 |
| CLEC4E | CCL15 | IGHG1 | MX2 | CCL2 |
| MMP9 | CCL16 | IGHM | MX1 | CCL20 |
| TMCC1 | CCL17 | IGHA1 | SAMHD1 | CCL21 |
| AC068580.4 | CCL18 | IGHV1-69-2 | ISG20 | CCL22 |
| CTSD | CCL19 | RAB31 | HLA-F | CCL23 |
| ARG2 | CCL2 | TRBC2 | IP6K2 | CCL24 |
| FPR1 | CCL20 | CDC42 | YTHDF2 | CCL25 |
| SLC16A3 | CCL21 | THBS1 | ZBP1 | CCL26 |
| JUNB | CCL22 | IGLL5 | LSM14A | CCL27 |
| DUSP1 | CCL23 | IGLC2 | CNOT7 | CCL28 |
| RDH12 | CCL24 | IGHV4-30-4 | IRF3 | CCL3 |
| SLC7A11 | CCL25 | IGHV3-43D | TYK2 | CCL3L1 |
| ASPRV1 | CCL26 | IGHV3-30-5 | IKBKE | CCL4 |
| S100A11 | CCL3 | IGHV3-30-3 | HLA-G | CCL4L1 |
| TIMP2 | CCL3L1 | IGHV1-8 | STAT1 | CCL5 |
| MXD1 | CCL4 | ARHGAP25 | IRF1 | CCL7 |
| CYP4F3 | CCL4L1 | IGLL1 | HLA-C | CCL8 |
| MAP1LC3B2 | CCL5 | GULP1 | PTPN11 | CKLF |
| MAP1LC3B | CCL7 | TRDC | ISG15 | CX3CL1 |
| YPEL3 | CCL8 | RHOG | USP18 | CXCL1 |
| CCR1 | CCR7 | PPARG | TREX1 | CXCL10 |
| FTL | CD300H | IGHV4-31 | IRF8 | CXCL11 |
| IL36G | CD74 | IGHV4-38-2 | IRF6 | CXCL12 |
| SLPI | CKLF | IGLC3 | IFIT3 | CXCL13 |
| RETNLB | CSF3R | ABCA1 | IRF9 | CXCL14 |
| CSTA | CX3CL1 | STAP1 | IFNA16 | CXCL16 |
| CD300LF | CXADR | XKR6 | IFNA4 | CXCL2 |
| FTH1 | CXCL1 | XKR7 | IFNA6 | CXCL3 |
| HACD4 | CXCL10 | XKR9 | TTLL12 | CXCL5 |
| MSRB1 | CXCL11 | IGHV2-70D | HLA-E | CXCL6 |
| IFITM1 | CXCL13 | IGHV3-66 | HLA-H | CXCL8 |
| IFITM2 | CXCL2 | IGHV4-61 | HLA-B | CXCL9 |
| IFITM3 | CXCL3 | IGHV1-58 | IFI27 | GPR15L |
| MMP8 | CXCL5 | IGHV5-51 | IRF2 | PF4 |
| S100A6 | CXCL6 | IGHV3-38 | PTPN1 | PF4V1 |
| CXCR2 | CXCL8 | IGHV3-35 | C9JQL5 | PPBP |
| IL1B | CXCL9 | IGHV4-28 | JAK1 | XCL1 |
| STK17B | CXCR1 | IGHV1-24 | IRF5 | XCL2 |
| | CXCR2 | IGHV3-20 | ADAR | |
| | DAPK2 | IGHV1-18 | BST2 | |

| | | |
|---|---|---|
| DNM1L | IGHV3-16 | RNASEL |
| DPEP1 | IGHV1-3 | EGR1 |
| DPP4 | ITGA2 | IFNA14 |
| EDN1 | MARCO | IFNA7 |
| EDN2 | CD300A | IFNA1 |
| EDN3 | VAMP7 | IFI35 |
| FCER1G | S4R3C0 | HSP90AB1 |
| GBF1 | HAVCR1 | MAVS |
| ITGA1 | RAC1 | MYD88 |
| ITGB2 | ARHGAP12 | IRF4 |
| JAML | BIN2 | CACTIN |
| LBP | APPL2 | IRF7 |
| LGALS3 | MFGE8 | RSAD2 |
| MCU | MEGF10 | MMP12 |
| MDK | FCGR1A | OASL |
| MOSPD2 | IGLC7 | IFNAR2 |
| NCKAP1L | XKR5 | PSMB8 |
| PDE4B | XKR4 | SP100 |
| PF4 | A0A0J9YWU9 | H0Y3Z8 |
| PF4V1 | GSN | ABCE1 |
| PIK3CD | IGHV4-4 | HLA-A |
| PIK3CG | IGHV1-2 | NLRC5 |
| PIKFYVE | C3 | WNT5A |
| PIP5K1C | ITGAM | METTL3 |
| PLA2G1B | IGHV5-10-1 | IFIT1 |
| PPBP | A0A0J9YY99 | IFIT2 |
| PPIA | ABCA7 | IFI6 |
| PPIB | SH3BP1 | IRAK1 |
| PREX1 | MSR1 | IFNA2 |
| RAC1 | IGLC1 | IFNA10 |
| RAC2 | ALOX15 | IFNA21 |
| RIPOR2 | ELMO1 | IFNA5 |
| S100A12 | IGHV1-69 | IFNA17 |
| S100A8 | NCKAP1L | IFNB1 |
| S100A9 | RHOBTB2 | IFITM1 |
| SAA1 | IGHV1-69D | UBE2K |
| SLIT2 | IGHV4-34 | FADD |
| SRP54 | RHOBTB1 | TRIM6 |
| SYK | IGHV3-64D | OAS2 |
| TGFB2 | IGLC6 | XAF1 |
| THBS4 | IGHV3-9 | MUL1 |
| TIRAP | IGHV3-7 | DCST1 |
| URS00000B7E30_9606 | IGHV3-33 | IFITM2 |
| VAV1 | IGHV3-30 | IFITM3 |
| VAV3 | IGHV3-53 | YTHDF3 |
| XCL1 | IGHV3-13 | OAS1 |
| XCL2 | IGHV3-23 | OAS3 |
| | IGHV3-48 | |
| | IGHV3-11 | |
| | IGHV1-46 | |
| | IGHV6-1 | |
| | IGHV3-21 | |
| | IGHV2-26 | |
| | IGHV3-73 | |
| | IGHV7-4-1 | |
| | A0A0J9YW62 | |
| | ITGB2 | |
| | FCER1G | |
| | IGHV1OR21-1 | |
| | GATA2 | |
| | ADGRB1 | |
| | IGHV3-15 | |
| | IGHV7-81 | |
| | DOCK1 | |
| | XKR8 | |
| | IGHV3-43 | |
| | IGHD | |
| | IGHA2 | |

| | | |
|---|---|---|
| | | IGHG4 |
| | | IGHG3 |
| | | IGHG2 |
| | | IGHE |
| | | RAC2 |
| | | IGHV1OR15-9 |
| | | IGHV2OR16-5 |
| | | IGHV3-72 |
| | | IGHV3-74 |
| | | IGHV4-30-2 |
| | | IGHV3-49 |
| | | IGHV1-45 |
| | | FCGR2B |
| | | IGHV3OR16-8 |
| | | IGHV3OR16-10 |
| | | IGHV3OR16-13 |
| | | IGHV3OR15-7 |
| | | IGHV1OR15-1 |
| | | IGHV3OR16-12 |
| | | IGHV4OR15-8 |
| | | ANO6 |
| | | F2RL1 |
| | | AIF1 |
| | | RHOH |
| | | CD36 |
| | | IGHV3-64 |
| **Exhaustion** | **Cytotoxicity** | **Treg** |
| TIGIT | KLRG1 | IL2RA |
| PDCD1 | KLRD1 | FOXP3 |
| CTLA4 | GZMK | |
| CD160 | GZMB | |
| KLRC1 | GZMA | |
| BTLA | GNLY | |
| HAVCR2 | PRF1 | |
| LAG3 | GZMM | |
| | NKG7 | |
| | TBX21 | |
| | ZEB2 | |
| | HOPX | |
| | GZMH | |
| | KLRK1 | |
| | IFNG | |
| | CCL3 | |
| | CST7 | |
| | ADGRG1 | |
| | IL32 | |
| | CRTAM | |
| | CX3CR1 | |
| | KLRC1 | |
| | FGFBP2 | |
| | FCGR3A | |
| | NCR3 | |
| | CCL4 | |

**Supplementary Table S4**

| Celltype | Abbreviation | Markers | Reference |
|---|---|---|---|
| Hepatocytes | Hepatocytes | ALB,TTR,APOA1 | 10.1186/s13059-020-02210-0,10.1016/j.cell.2020.11.041,10.1016/j.cell.2020.11.041 |
| Cholangiocytes | Cholangiocytes | KRT19,KRT7,CFTR | 10.1186/s13059-020-02210-0,10.1186/s13059-020-02210-0,10.1186/s13059-020-02210-0 |
| Endothelial cells | ECs | PECAM1,VWF,CDH5 | 10.7150/thno.54917,10.1158/1078-0432.CCR-19-3231,10.1038/s41556-019-0446-7 |
| Hepatic stellate cells | HepSCs | PDGFRB,ACTA2,RGS5 | 10.1172/JCI146987,10.1016/j.cell.2020.11.041,10.1016/j.cell.2020.11.041 |
| Proliferating cells | ProliferatingCells | MKI67,TOP2A,STMN1 | 10.1158/1078-0432.CCR-19-3231,10.1681/ASN.2019080832,10.1038/s41467-019-14256-1 |
| B cells | BCells | CD79A,MS4A1,CD19 | 10.1038/s41467-021-21795-z,10.1038/s41467-021-21795-z,10.1038/s41422-020-0378-6 |
| Plasma cells | PlasmaCells | JCHAIN,CD79A,MZB1 | 10.1038/s41467-021-21795-z,10.1038/s41467-021-21795-z,10.1038/s41467-019-14256-1 |
| T and NK cells | TandNK | CD3D,CD3E,NKG7 | 10.1038/s41467-021-21795-z,10.1016/j.immuni.2019.09.008,10.1530/ERC-22-0325 |
| Neutrophils | Neutrophils | FCGR3B,S100A9,S100A8 | 10.1038/s41467-021-22801-0,10.1038/s41467-021-22801-0,10.1038/s41467-021-22801-0 |
| Mast cells | MastCells | TPSAB1,TPSB2,CPA3 | 10.1158/1078-0432.CCR-19-3231,10.1038/s41467-021-22801-0,10.1016/j.immuni.2019.09.008 |
| Mononuclear phagocytes | MPs | CD14,CSF1R,HLA-DRA | 10.1038/s41586-019-1631-3,10.1038/s41586-019-1631-3,10.1038/s41586-019-1631-3 |
| Plasmacytoid dendritic cells | pDCs | IL3RA,CLEC4C,LILRA4 | 10.1158/1078-0432.CCR-19-3231,10.1158/1078-0432.CCR-19-3231,10.1038/s41593-020-00789-y |

| Celltype | Abbreviation | Markers | Reference |
|---|---|---|---|
| Proliferating cells | ProliferatingCells | MKI67,TOP2A,STMN1 | 10.1158/1078-0432.CCR-19-3231,10.1681/ASN.2019080832,10.1038/s41467-019-14256-1 |
| Macrophages | Macrophages | CD14,C1QA,CSF1R | 10.1158/1078-0432.CCR-19-3231,10.1016/j.immuni.2019.09.008,10.1038/s41556-019-0446-7 |
| Monocytes | Monocytes | LYZ,FCN1,VCAN | 10.1038/s41467-019-11049-4,10.1038/s41467-021-22801-0,10.1038/s41467-021-22801-0 |
| Mature dendritic cells | MatureDCs | LAMP3,CCR7,CD83 | 10.1158/2159-8290.CD-19-0138,10.1038/s41591-021-01323-8,10.1158/2159-8290.CD-19-0138 |
| Conventional type 1 dendritic cells | cDC1 | CLEC9A,XCR1,IRF8 | 10.1038/s41586-019-1652-y,10.1038/s41586-019-1652-y,10.1038/s41422-020-0374-x |
| Conventional type 2 dendritic cells | cDC2 | CD1C,CLEC10A,FCER1A | 10.1038/s41586-019-1652-y,10.1038/s41586-019-1652-y,10.1038/s41593-020-00789-y |
| Kupffer cells | KCs | MARCO,CD5L,VCAM1 | 10.1016/j.cell.2020.11.041,10.1016/j.cell.2020.10.048,10.1038/s41586-019-1631-3 |

| Celltype | Abbreviation | Markers | Reference |
|---|---|---|---|
| Group 3 innate lymphoid cells | ILC3 | IL1R1,IL23R,KIT | 10.1038/s41590-019-0425-y,10.1038/s41590-019-0425-y,10.1038/s41586-021-03852-1 |
| NK T cells | NKT | NKG7,CD3D,GNLY | 10.7150/thno.48201,10.7150/thno.48201,10.7150/thno.48201 |
| Natural killer cells | NK | NKG7,GNLY,NCAM1 | 10.1038/s41392-020-00248-x,10.1038/s41467-021-21795-z,10.1126/science.aad0501 |
| CD4+ naive T cells | CD4NaiveT | CCR7,SELL,LEF1 | 10.1038/s41467-021-22164- |
| CD4+ memory T cells | CD4Tmem | CD4,IL7R,CD40LG | 10.1182/blood-2018-08-862292,10.1016/j.immuni.2020.12.011,10.1038/s41467-022-33170-7 |
| CD4+ regulatory T cells | CD4Treg | CTLA4,FOXP3,IL2RA | 10.1038/s41421-020-0157-z,10.1038/s41421-020-0157-z,10.1038/s41421-020-0157-z |
| CD8+ mucosal-associated invariant T cells | CD8MAIT | KLRB1,SLC4A10,NCR3 | 10.1016/j.cell.2021.05.013,10.1186/s13059-019-1906-x,10.1038/s41392-020-00263-y |
| CD8+ effector T cells | CD8Teff | CD8A,GZMK,CD3D | 10.1038/s41467-021-22801-0,10.1038/s41591-019-0590-4,10.1016/j.cell.2020.06.001 |
